# Hypoxia Sensing in Resident Cardiac Macrophages Regulates Monocyte Fate Specification following Ischemic Heart Injury

**DOI:** 10.1101/2022.08.04.502542

**Authors:** Farid F. Kadyrov, Andrew L. Koenig, Junedh M. Amrute, Hao Dun, Wenjun Li, Carla J. Weinheimer, Jessica M. Nigro, Attila Kovacs, Andrea L. Bredemeyer, Lulu Lai, Benjamin J. Kopecky, Vinay Penna, Daniel Kreisel, Kory J. Lavine

## Abstract

Myocardial infarction initiates cardiac remodeling and is central to heart failure pathogenesis. Following myocardial ischemia reperfusion injury, monocytes enter the heart and differentiate into diverse subpopulations of macrophages. The mechanisms and dynamics of monocyte differentiation within this context are unknown. We investigated the role of macrophage hypoxia sensing on monocyte differentiation following reperfused myocardial infarction. We show that deletion of *Hif1α*, a hypoxia response transcription factor, in resident cardiac macrophages led to increased remodeling and overrepresentation of a macrophage subset marked by arginase 1 (*Arg1*) expression. Arg1^+^ macrophages displayed an inflammatory gene signature and were predicted to represent an intermediate state within the monocyte differentiation cascade. Lineage tracing of Arg1^+^ macrophages revealed the existence of a monocyte differentiation trajectory consisting of multiple transcriptionally distinct macrophage states. We further showed that deletion of *Hif1α* in resident cardiac macrophages resulted in arrested progression through this trajectory and accumulation of an inflammatory intermediate state marked by persistent *Arg1* expression. Collectively, our findings unveil distinct trajectories of monocyte differentiation and identify hypoxia sensing as an important determinant of monocyte differentiation following myocardial infarction.

## Introduction

Myocardial infarction (MI) and other forms of ischemic cardiac injury initiate adverse left ventricular (LV) remodeling and are central to the pathogenesis of heart failure^1,2^. Heart failure continues to represent a serious and life threatening complication of MI despite the availability of effective medications that limit adverse LV remodeling^1–3^. Activation of innate immune responses and the resultant cardiac inflammation which occurs after MI may represent a promising therapeutic target to limit LV remodeling and heart failure^4^.

Macrophages represent the primary immune cell found in the heart under homeostasis. Cardiac macrophages are complex, arising from multiple lineages. The mouse and human heart contain two functionally distinct resident macrophage populations distinguished by C-C chemokine receptor 2 (CCR2) expression: embryonic-derived tissue resident CCR2^−^ macrophages that promote tissue repair and suppress inflammation, and monocyte-derived CCR2^+^ macrophages that initiate inflammatory responses^5–8^. Following MI, large numbers of Ly6C^hi^CCR2^+^ monocytes are recruited to and infiltrate the heart and differentiate into diverse macrophage populations that are traditionally viewed as having pro-inflammatory properties^9,10^.

Prior studies have elucidated that manipulating cardiac macrophage composition and interfering with monocyte recruitment has dramatic impacts on outcomes following MI^5,6,11–13^. Depletion of CCR2^+^ macrophages prior to MI is sufficient to reduce neutrophil and monocyte infiltration, suppress myocardial chemokine/cytokine production, and preserve LV systolic function. Conversely, removing CCR2^−^ tissue resident macrophages leads to increased peripheral leukocyte recruitment and accelerated LV remodeling. Consistent with a deleterious role for infiltrating leukocytes, inhibition of monocyte recruitment has beneficial effects in mouse models of MI^5,12–15^.

Recent single cell RNA sequencing (scRNAseq) studies have indicated that monocytes recruited to the injured heart differentiate into multiple distinct macrophage and dendritic-like cell subsets^5^. The mechanistic basis and dynamics by which monocytes differentiate into these populations are unknown. We hypothesized that environmental cues present within the infarcted heart instruct monocyte differentiation decisions. These cues may either act directly on infiltrating monocytes or indirectly through non-cell autonomous mechanisms. The existence of non-cell autonomous pathways is supported by the observation that depletion of cardiac resident macrophages prior to MI is sufficient to alter the relative abundance of various monocyte-derived macrophage subsets^5^.

Here, we explored hypoxia sensing as an environmental cue that may influence monocyte differentiation following MI using mouse models of myocardial ischemia reperfusion injury. Hypoxia inducible factor-1 (HIF-1) is a transcription factor that regulates the cellular response to hypoxia^16–18^. We conditionally deleted *Hif1α* in either all monocytes and macrophages, recruited monocytes and macrophages, or resident macrophages. We demonstrate that *Hif1α* deletion specifically in resident macrophages accelerated LV remodeling and led to an expansion of a subpopulation of monocyte-derived macrophages marked by arginase 1 (*Arg1*) expression. Arg1^+^ macrophages represented a population that was specified early during monocyte differentiation and displayed an inflammatory gene signature. Lineage tracing studies demonstrated that Arg1^+^ macrophages differentiate into multiple distinct macrophage subsets that included Gdf15^+,^ Trem2^+^, MHC-II^hi^, and resident-like macrophages. *Hif1α* deletion in resident macrophages arrested progression through this lineage and resulted in accumulation of pro-inflammatory macrophages with persistent *Arg1* expression. These findings identify distinct lineages of monocyte-derived macrophages within the infarcted heart and identify hypoxia sensing in resident cardiac macrophages as a non-cell autonomous mediator of monocyte differentiation.

## Results

### Deletion of *Hif1α* in monocytes and macrophages results in accelerated LV remodeling following MI

To assess the functional requirement for hypoxia sensing in monocytes and macrophages following MI, we subjected littermate control and *Hif1α^flox/flox^ Cx3cr1^ERT2Cre^* mice to closed-chest ischemia reperfusion injury (I/R) (90 minutes of ischemia) to simulate reperfused MI^19,20^. Tamoxifen chow was given throughout the entirety of the experiment to induce *Hif1α* deletion in all monocytes and macrophages (macHif1aKO) (**Fig. 1a, Extended Data Fig. 1a**). Echocardiography performed 4 weeks after I/R revealed that macHif1aKO mice had increased akinetic LV area, reduced LV ejection fraction, and increased LV diastolic and systolic volumes compared to controls (**Fig. 1b-c**). Trichrome and WGA staining performed 4 weeks after I/R revealed increased infarct size and cardiomyocyte hypertrophy in macHif1aKO mice (**Fig. 1d-g**). TTC staining performed within 4 days after I/R showed no significant difference in initial infarct area between control and macHif1aKO mice, indicating that increased final infarct size in macHif1aKO mice is due to infarct expansion rather than an increase in the initial extent of injury (**Fig. 1f**).

**Fig. 1.**
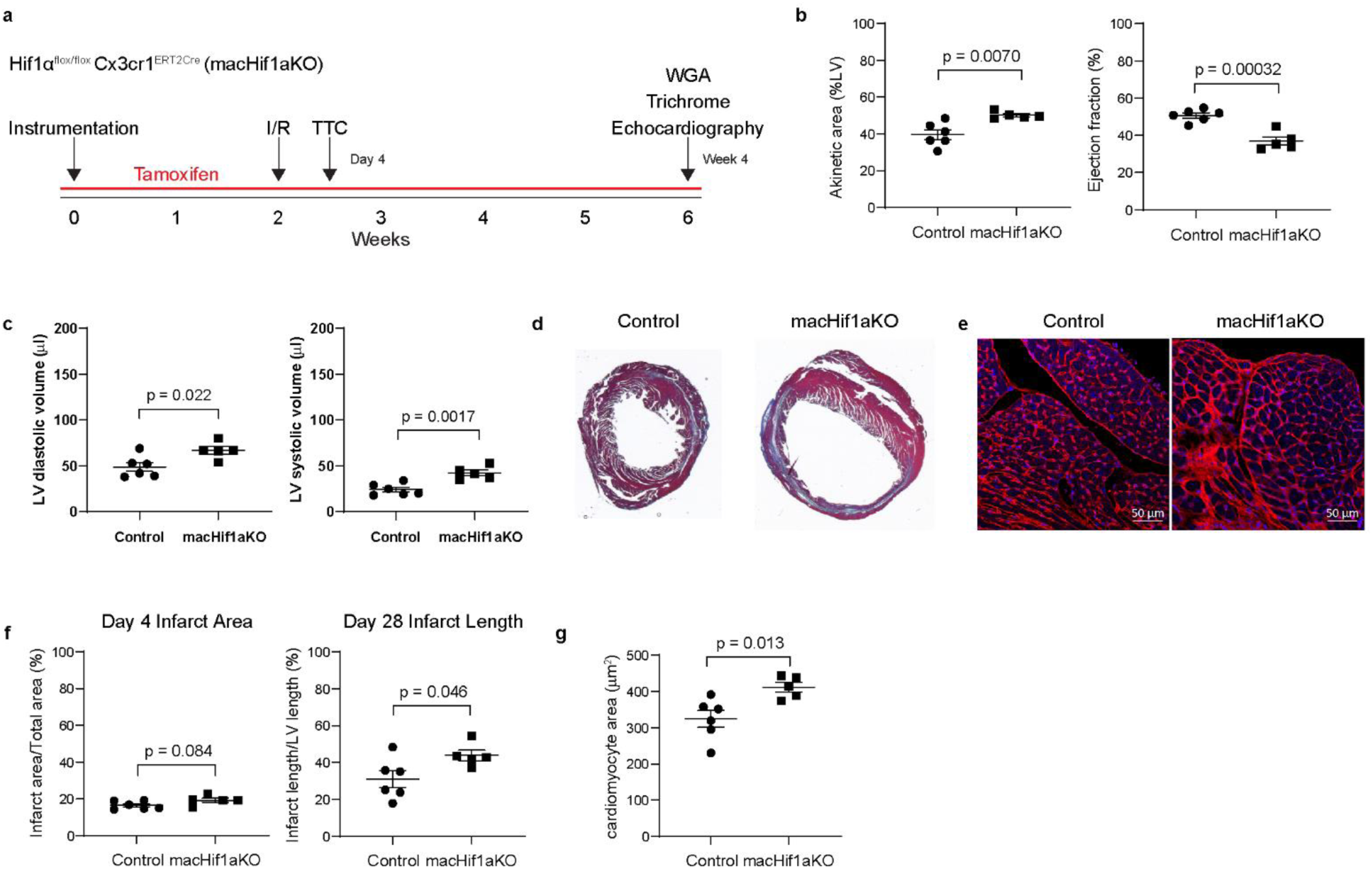
Hif1α deletion in monocytes and macrophages leads to accelerated LV remodeling and reduced cardiac function after MI. **a,** Experimental outline of the ischemia reperfusion injury (I/R) mouse model of MI in macHif1aKO mice. Tamoxifen chow was given throughout the course of the experiment. Closed chest I/R was performed 2 weeks after instrumentation, and echocardiography was performed 4 weeks after I/R. Tissues were collected 4 days and 4 weeks after I/R for analysis. **b,** Echocardiography quantification 4 weeks after I/R of akinetic area as a percentage of the left ventricle and ejection fraction between Control and macHif1aKO mice. **c,** Left ventricular (LV) diastolic volume, and LV systolic volume between Control and macHif1aKO mice. **d,** Representative trichrome staining of Control and macHif1aKO hearts 4 weeks after I/R. **e,** Representative WGA staining of the border zone of the infarct of Control and macHif1aKO hearts 4 weeks after I/R. **f,** Quantification of TTC staining 4 days after I/R to determine initial infarct area (displayed as a percentage of infarct area/total area) and quantification of trichrome staining in d (displayed as the length of the infarct as a percentage of the total length of the left ventricle) between Control and macHif1aKO mice. **g,** Quantification of the average cardiomyocyte area determined from WGA staining in e in Control and macHif1aKO hearts. P-values are determined using a two tailed t-test assuming equal variance.

In the context of MI and heart failure, *Hif1α* is thought to contribute to expansion of the coronary vasculature within the infarct and border zones^21^. Immunostaining (IHC) for CD34 (microvasculature marker) and smooth muscle actin (α-SMA, large blood vessel marker) did not reveal any reduction in vascular density in macHif1aKO hearts following I/R (**Extended Data Fig. 1b**-d)^22^. To assess whether *Hif1α* deletion in monocytes and macrophages alters the abundance of the myeloid cells that infiltrate the heart after MI^23^, we performed IHC and flow cytometry analysis (FACS) of control and macHif1aKO mice 4-5 days after I/R. We observed no differences in the total number of monocytes or macrophages between conditions and a marginal reduction in neutrophils in macHif1aKO hearts (**Extended Data Fig. 1e**-i).

### *Hif1α* deletion in monocytes and macrophages results in a shift in monocyte fate specification following MI

Previous scRNAseq studies have revealed that altering the identity of macrophages present within the heart following MI has profound effects on LV remodeling^5,14^. To evaluate if *Hif1α* deletion in monocytes and macrophages impacts the differentiation of infiltrating monocytes following I/R, we utilized scRNAseq as a tool to identify transcriptionally distinct populations of monocytes and their progeny (macrophages and dendritic-like cells) in control and macHif1aKO mice. scRNAseq was performed on cardiac monocytes, macrophages, and dendritic-like cells isolated by FACS 5 days after I/R (**Fig. 2a-b, Extended Data Fig. 2a**-b). To consistently annotate cell populations with high confidence, scRNAseq data generated from control and macHif1aKO mice was mapped using label transfer^24^ to a reference data set of myeloid cells isolated from the mouse heart 3, 7, 14, and 28 days after MI^25^ (**Extended Data Fig. 2c**). Reference mapping performance was quantified and validated by calculating mapping scores and plotting transcriptional signatures of reference cell populations in the query dataset (**Fig. 2c, Extended Data Fig. 2d**-h).

**Fig. 2.**
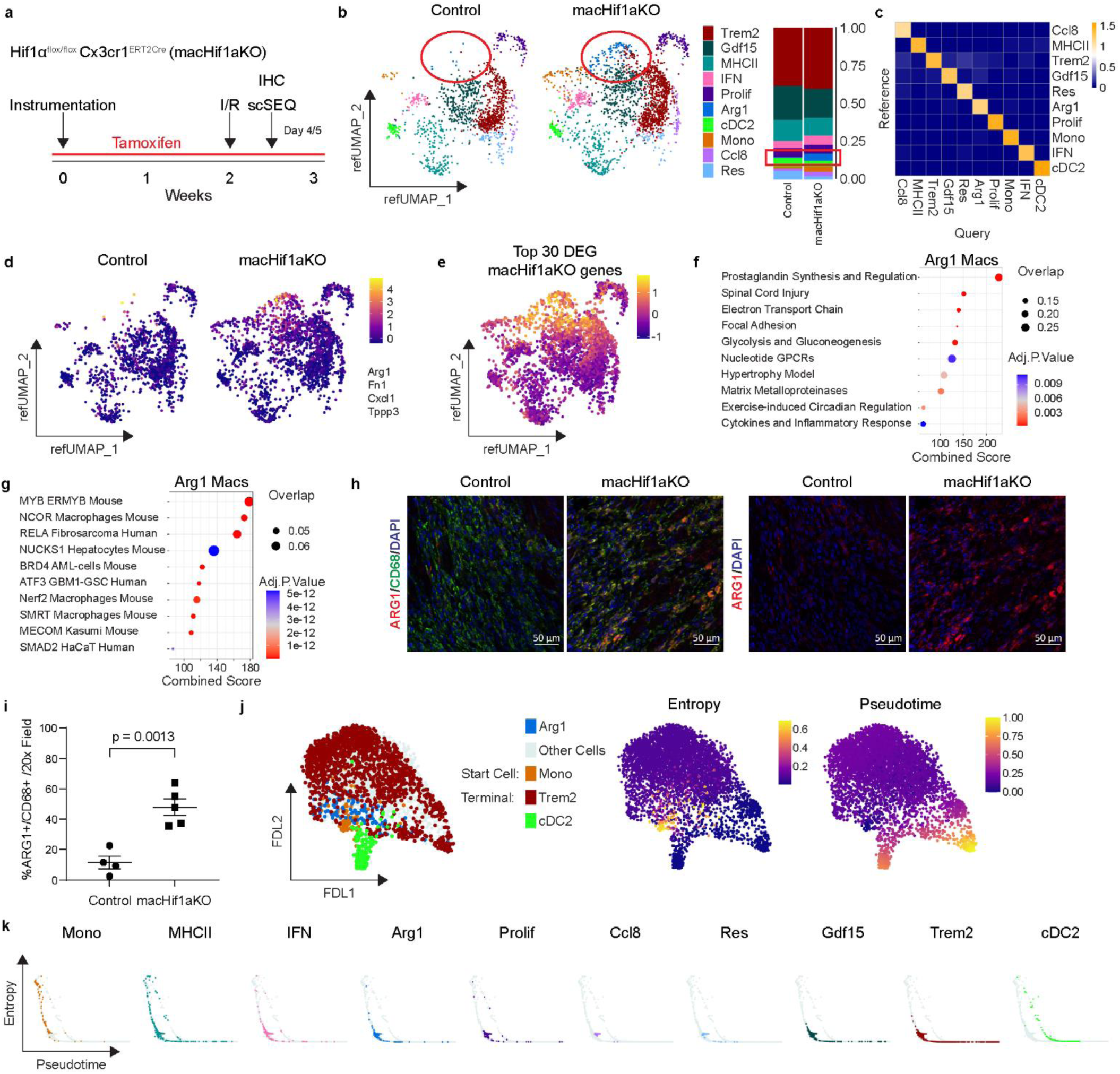
Single cell RNA sequencing of monocytes, macrophages, and dendritic-like cells 5 days after MI reveals overrepresentation of Arg1+ macrophages in macHif1aKO mice. **a,** Experimental outline of the I/R mouse model of MI in macHif1aKO mice. Tissues were collected 4 days after I/R for IHC and 5 days after I/R for scSEQ. **b,** Annotated reference UMAP of scSEQ data, and stacked bar graph representing the proportion of each cell population split between Control and MacHif1aKO mice. Data was mapped to a reference MI data set of monocytes/macrophages/dendritic-like cells 3, 7, 13, and 28 days after I/R. Red circles and box highlight Arg1^+^ macrophages in each condition. **c,** Heat map of mapping scores for each cell population in the query (Control and macHif1a KO scSEQ) plotted against the reference MI data set. **d,** Z-score profile of Arg1^+^ macrophages (*Arg1*, *Fn1*, *Cxcl1*, *Tpp3*) plotted on a reference UMAP and split between conditions. **e,** Z-score profile of the top 30 DEG upregulated in macHif1aKO mice plotted on a reference UMAP. **f,** Dot plot of the top 10 upregulated pathways (Enrichr Wikipathways 2019 Mouse) in Arg1^+^ macrophages. **g,** Dot plot of the top 10 ChEA 2016 EnrichR transcription factors upregulated in Arg1^+^ macrophages. **h,** 20x confocal images of ARG1 (red), CD68 (green), and DAPI (blue) IHC staining within the infarct zone 4 days after I/R of Control and macHif1aKO mice. **i,** Quantification of IHC in h, displayed as the percentage of CD68^+^ cells that are ARG1^+^CD68^+^. **j,** FDL representation of Palantir analysis displaying the starting cells (Monocytes, orange), predicted terminal states (Trem2, red; cDC2; green), Arg1 cells (blue) and all other cells (gray). Entropy and pseudotime scores from Palantir plotted on FDL. **k,** Plots of entropy vs pseudotime from Palantir for each subpopulation. P-values are determined using a two tailed t-test assuming equal variance.

Composition analysis of monocytes, macrophages, and dendritic-like cells in control and macHif1aKO mice 5 days after I/R demonstrated a overrepresentation of a macrophage population marked by *Arg1* mRNA expression in macHif1aKO mice (**Fig. 2a-b**), suggesting that hypoxia sensing in monocytes and/or macrophages may influence the differentiation of infiltrating monocytes. Arg1^+^ macrophages had a unique transcriptional signature with significant enrichment in *Arg1*, *Fn1*, *Cxcl1*, and *Tppp3* mRNA expression (**Fig. 2d**). Differential gene expression analysis comparing control and macHif1aKO conditions revealed that many of the genes increased in macHif1aKO monocytes, macrophages, and dendritic-like cells were selectively expressed by Arg1^+^ macrophages (**Fig. 2e**). Given the increased abundance of Arg1^+^ macrophages and mapping of differentially expressed genes to this population, we further examined the transcriptional profile of Arg1^+^ macrophages using pathway analysis. Arg1^+^ macrophages were characterized by the expression of genes involved in prostaglandin synthesis, metabolic regulation (electron transport chain, glycolysis, gluconeogenesis) and cytokine and inflammatory response pathways (**Fig. 2f**). Transcription factor enrichment predicted that marker genes of Arg1^+^ macrophages were regulated by transcription factors involved in the development and function of inflammatory macrophages (MYB, NCOR, RELA, BRD4, ATF3, SMRT, NRF2) (**Fig. 2g**)^26–31^. IHC co-staining for ARG1 and CD68 (macrophage marker) confirmed the overrepresentation of Arg1^+^ macrophages within the infarct of macHif1aKO mice 5 days after I/R (**Fig. 2h-i**).

To ascertain the differential trajectory of monocytes, macrophages, and dendritic-like cells in control and macHif1aKO hearts we used Palantir trajectory analysis^32^. Cell cycle regression was performed prior to trajectory analysis^33^ (**Extended Data Fig. 3a**). Monocytes were selected as the least differentiated cell population. Palantir predicted Trem2^+^ macrophages and cDC2s as terminal populations (low entropy and high pseudotime values) (**Fig. 2j, Extended Data Fig. 3b**-d). As anticipated, monocytes displayed high entropy and low pseudotime values. Intriguingly, Arg1^+^ macrophages displayed entropy and pseudotime values in between monocytes and terminal populations, suggesting that they may represent an intermediate in monocyte differentiation (**Fig. 2j-k**). These data support the concept that hypoxia sensing in monocytes and/or macrophages may influence steps in monocyte fate specification.

### Arg1^+^ macrophages are monocyte-derived, infiltrate early after ischemic injury, and are specified prior to myocardial extravasation

We next validated the above predictions made from analysis of our scRNAseq data. To determine if Arg1^+^ macrophages are monocyte derived, we utilized a monocyte fate labeling strategy. To label monocytes and their progeny, we established a tamoxifen regimen where a single dose of 60mg/kg administered to *Ccr2^ERT2Cre^ Hif1α^flox/flox^ Rosa26^lsl-tdT^* (monoTDT) mice labels peripheral monocytes and not cardiac macrophages. We subjected monoTDT mice to closed chest I/R and assessed tdTomato (TDT) expression 5 days after I/R (**Fig. 3a**). Mice were treated with a single dose of tamoxifen (60mg/kg) 24 hours prior to I/R. The overwhelming majority of Arg1^+^ macrophages were TDT^+^ indicating that Arg1^+^ macrophages are derived from monocytes. The same observation was made with *Ccr2^ERT2Cre^ Rosa26^lsl-tdT^*mice 3 days after I/R (**Fig. 3b**).

**Fig. 3.**
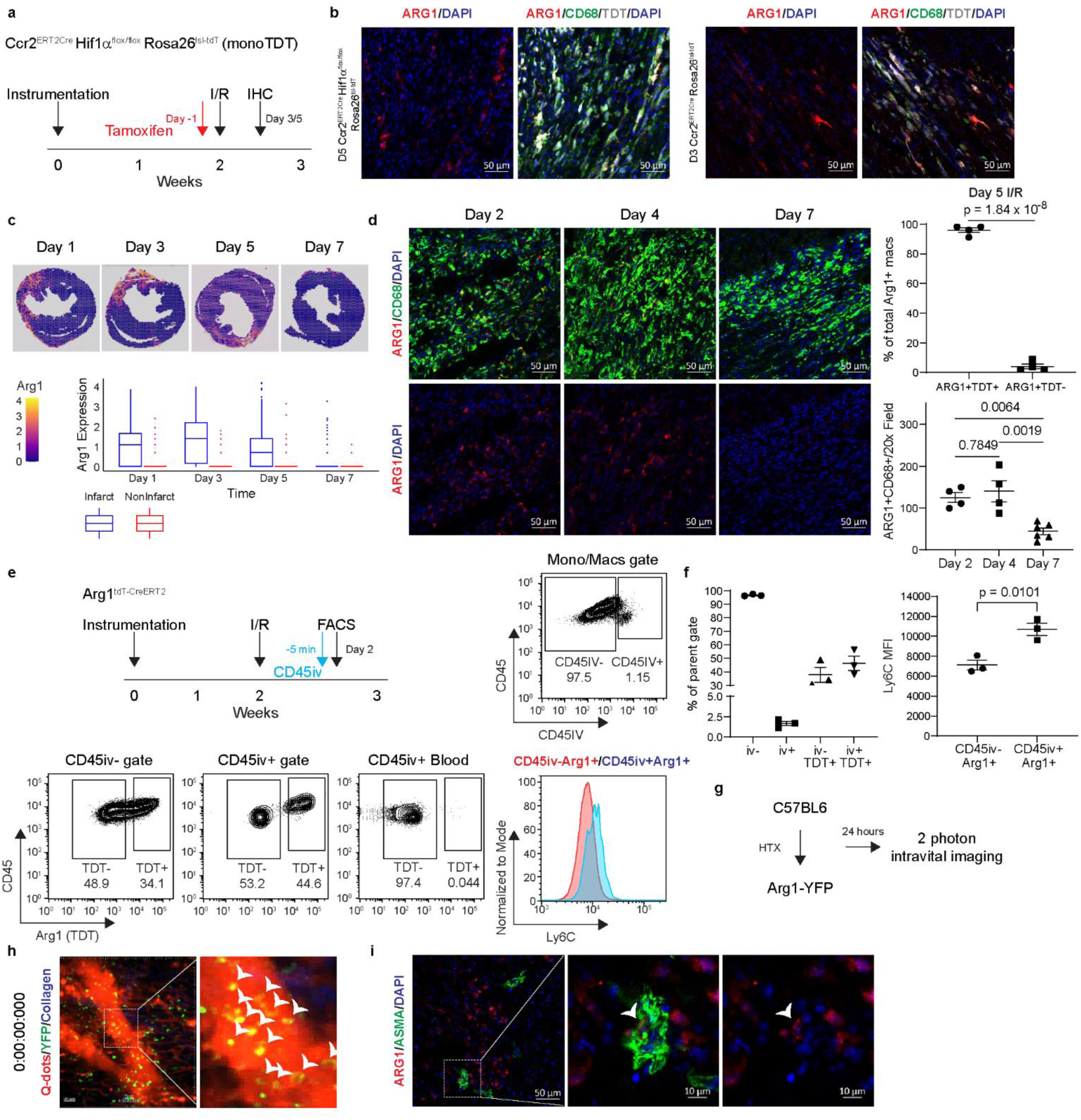
Spatiotemporal dynamics of Arg1+ macrophages after ischemic heart injury. **a,** Schematic of the mouse model of MI in monoTDT mice. 60 mg/kg of tamoxifen was IP injected 1 day prior to I/R. Tissues were collected 3/5 days after I/R for IHC. **b,** Representative 20x confocal image of ARG1 (red), CD68 (green), TDT (white) and DAPI (blue) IHC within the infarcts of monoTDT mice. Quantification displayed as the percentage of ARG1^+^CD68^+^ macrophages that are ARG1^+^TDT^+^ or ARG1^+^TDT^−^. **c,** *Arg1* RNA expression of a spatial transcriptomics dataset 1/3/5/7 days after MI (GSE165857). Expression between infarcted and non-infarcted regions are displayed as a box plot. **d,** Representative 20x confocal images of ARG1 (red), CD68 (green) and DAPI (blue) IHC within the infarct 2, 4, and 7 days after I/R. Quantification displayed as the total ARG1^+^CD68^+^ cells per 20x field. **e,** Schematic and representative FACS of *Arg1^tdT-CreERT^*^2^ mice 2 days after I/R. Mice were injected with a CD45 antibody 5 minutes prior to tissue collection. Histogram shows representative Ly6C staining between CD45iv^−^Arg1^+^ and CD45iv^+^Arg1^+^ cells. **f,** Quantification of populations of interest from e, and of Ly6C mean fluorescence intensity (MFI) between CD45iv^−^Arg1^+^ and CD45iv^+^Arg1^+^ cells. **g** Schematic of intravital 2-photon microscopy on coronary veins in *Arg1^YFP^* mice. C57BL6 hearts were syngeneically transplanted into *Arg1^YFP^* mice and 24 hours later were imaged for a duration of 15 minutes. **h,** Representative 2-photon image of syngeneic transplant *Arg1^YFP^* mice. Q-dots (red) were injected IV prior to imaging to visualize blood vessels. Collagen is in blue. Intravascular Arg1^+^ cells (green) co-stained with Q-dots are highlighted by arrows. **i,** 20x confocal image of IHC co-staining of ARG1 (red) and α-SMA+ vessels (green) 2 days after I/R highlighted by white arrow. P-values are determined using a two tailed t-test assuming equal variance or ordinary one-way ANOVA for multiple comparisons.

We next investigated the spatiotemporal dynamics of Arg1^+^ macrophages. A published spatial transcriptomics data set of mouse hearts 1, 3, 5, and 7 days after MI was used to gain insights into Arg1^+^ macrophage specification (GSE165857)^34^. RNA expression of *Arg1* was high within the infarct 1, 3, 5 days after MI, lower 7 days after MI, and largely absent in non-infarcted areas (**Fig. 3c, Supplementary Figure 1**). Consistent with this finding, IHC for ARG1 and CD68 in hearts subjected to closed chest I/R demonstrated Arg1^+^ macrophages present within the infarct 2 and 4 days after I/R, but largely gone 7 days after I/R (**Fig. 3d**).

Trajectory analysis predicted that Arg1^+^ macrophages represented an intermediate in monocyte differentiation. To investigate where Arg1^+^ macrophage specification is initiated, we employed a FACS-based approach to differentially label intravascular versus extravascular leukocytes. *Arg1^tdT-CreERT^*^2^ reporter mice^35^ were analyzed 2 days after MI, a time point where monocytes are robustly recruited to the heart^36^. A fluorescently conjugated CD45 antibody was injected intravascularly into *Arg1^tdT-CreERT^*^2^ reporter mice 5 minutes prior to tissue harvest, which has previously been shown to differentiate intravascular (CD45iv^+^) leukocytes from extravascular (CD45iv^−^) leukocytes^37^. We detected Arg1^+^ macrophages within both the extravascular (CD45iv^−^) and intravascular (CD45iv^+^) compartments. Conversely, negligible TDT signal was observed in peripheral monocytes, indicating that Arg1^+^ macrophages are specified within the vasculature of the heart (**Fig. 3e-f, Extended Data** Fig. 4). Consistent with the concept that intravascular and extravascular Arg1^+^ macrophages reflected different stages of monocyte/macrophage differentiation, mean fluorescence intensity of Ly6C staining was higher in Arg1^+^CD45iv^+^ cells compared to Arg1^+^CD45iv^−^ cells (**Fig. 3f**). Similar results were found with a second reporter line, *Arg1^YFP^* mice (**Supplementary Figure 2**).

To verify that Arg1^+^ macrophages are present within the intravascular compartment using an orthogonal approach, we performed intravital 2-photon microscopy to visualize Arg1^+^ macrophages in real time. We leveraged a previously established syngeneic heart transplant model of ischemia reperfusion injury in combination with *Arg1* reporter mice^5,38^. C57BL6 donor hearts were harvested, placed on ice for 60 minutes, and then transplanted into *Arg1^YFP^* mice in the cervical position (**Fig. 3g**). Fluorescently labeled Qdots were injected IV and donor hearts were imaged 24 hours after heart transplantation. We observed YFP^+^ cells within the intravascular and extravascular compartments, suggesting that Arg1^+^ macrophage specification is initiated prior to extravasation into the myocardium (**Fig. 3h**). ARG1 and α-SMA IHC confirmed Arg1^+^ cells within α-SMA+ vessels in control mice 2 days after I/R (**Fig. 3i**). Overall, these results reveal that Arg1^+^ macrophages are monocyte-derived and are specified early after monocyte recruitment to the ischemic heart.

### Conditional deletion of *Hif1α* in cardiac resident macrophages increases the abundance of Arg1^+^ macrophages and contributes to adverse outcomes following MI

We next investigated whether hypoxia sensing regulates Arg1^+^ macrophage specification in a cell intrinsic or extrinsic manner. To delete *Hif1α* exclusively in infiltrating monocytes and their progeny (monoHif1aKO), *Hif1α^flox/flox^ Ccr2^ERT2Cre^* mice received a single dose of tamoxifen (60mg/kg, IP) 24 hours prior to MI (**Fig. 4a**). To delete *Hif1α* exclusively in resident macrophages (resHif1aKO), we pulsed *Hif1α^flox/flox^ Cx3cr1^ERT2Cre^* mice with tamoxifen (60 mg/kg daily for 5 days) immediately after instrumentation to label all monocytes and macrophages. Mice then underwent a chase period of 9 days prior to MI (**Fig. 4a**). During this time monocytes are replenished from hematopoietic progenitors and only cardiac resident macrophages remain labeled^5^. IHC staining for CD68 and TDT within the infarct, border, and remote zones confirmed that these macrophage subsets occupy distinct locations within the infarcted heart (**Supplementary Figure 3**).

**Fig. 4.**
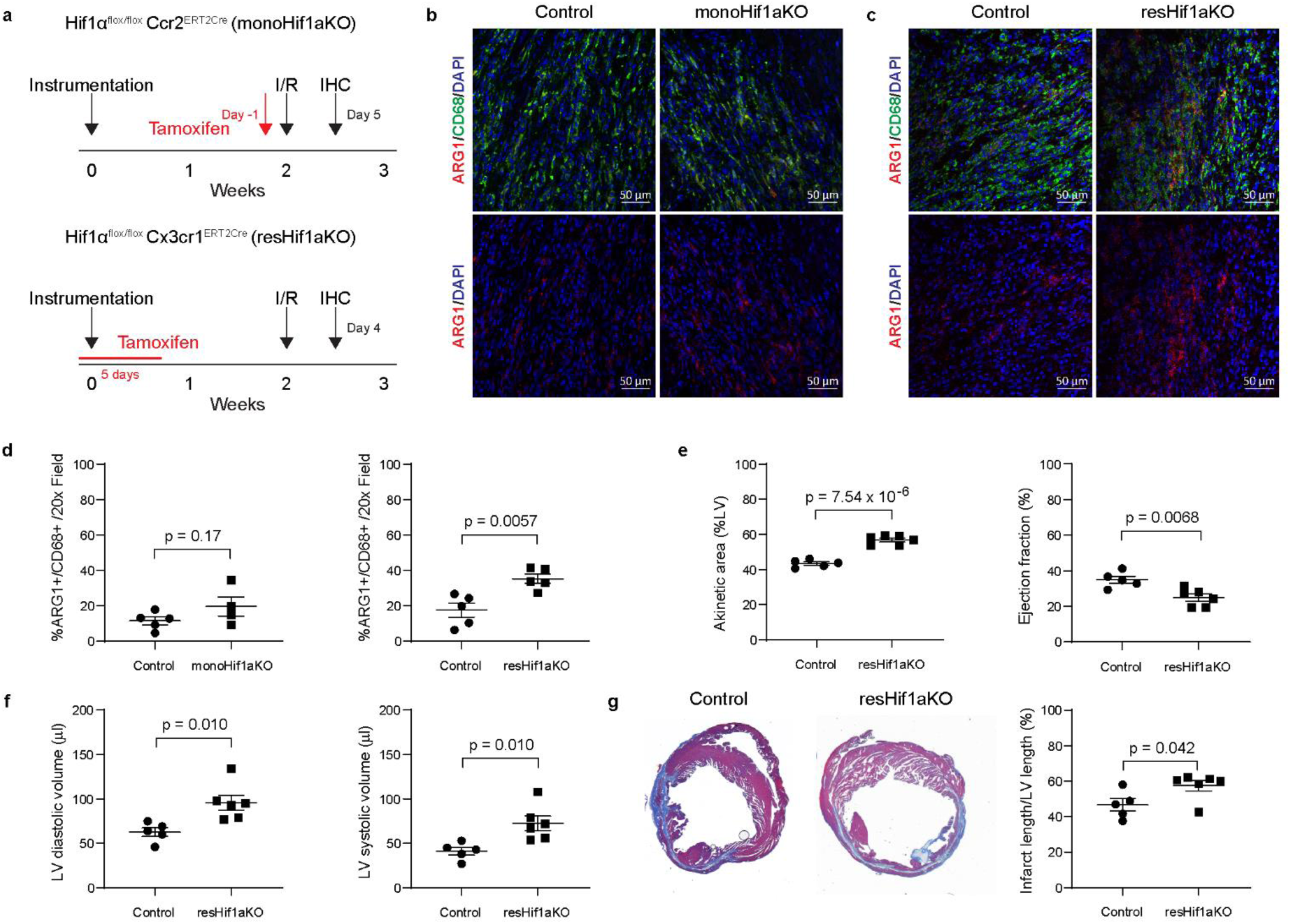
*Hif1α* deletion in cardiac resident macrophages is sufficient to increase the abundance of Arg1+ macrophages and accelerate LV remodeling after MI. **a,** Experimental outline of the I/R mouse model of MI in monoHif1aKO and resHif1aKO mice. Closed chest I/R was performed 2 weeks after instrumentation, and tissues were collected 4/5 days after I/R for analysis. 60 mg/kg of tamoxifen was IP injected 1 day prior to I/R in monoHif1aKO mice to induce *Hif1α* knockout in monocytes and recruited macrophages. 60 mg/kg of tamoxifen was IP injected for 5 days followed by a chase period to induce *Hif1α* knockout in resident macrophages. **b,** Representative 20x confocal images of IHC staining in monoHif1aKO and **c,** resHif1aKO mice. ARG1 is in red, CD68 is in green, DAPI is in blue. **d,** Quantification of IHC in b and c respectively. Data is displayed the percentage of ARG1^+^CD68^+^ cells out of total CD68^+^ cells. **e,** Echocardiography quantification 4 weeks after I/R of akinetic area as a percentage of the LV and ejection fraction between Control and resHif1aKO mice. **f,** LV diastolic and systolic volume between Control and resHif1aKO mice 4 weeks after I/R. **g,** Representative trichrome staining and quantification (displayed as the length of the infarct as a percentage of the total length of the left ventricle) of Control and resHif1aKO hearts 4 weeks after I/R. P-values are determined using a two tailed t-test assuming equal variance.

To delineate whether deleting *Hif1α* in infiltrating monocytes and/or cardiac resident macrophages contributed to expansion of Arg1^+^ macrophages, we quantified Arg1^+^ macrophages by IHC 4-5 days after I/R in monoHif1aKO and resHif1aKO hearts. monoHif1aKO hearts did not display an increase in Arg1^+^ macrophages compared to control (**Fig. 4b,d**). Conversely, resHif1aKO displayed an increase in the abundance of Arg1^+^ macrophages within the infarct (**Fig. 4c,d**). This data indicates that *Hif1α* in cardiac resident macrophages influences the specification of Arg1^+^ macrophages through a non-cell autonomous mechanism.

We next examined whether deletion of *Hif1α* in cardiac resident macrophages contributes to LV remodeling. Echocardiography performed 4 weeks post-I/R revealed that resHif1aKO mice had increased akinetic LV area, reduced LV ejection fraction, and increased LV diastolic and systolic volume compared to controls (**Fig. 4 e,f**). Trichrome staining 4 weeks after I/R revealed increased infarct size in resHif1aKO mice compared to controls (**Fig. 4g**). WGA staining did not show a significant increase in cardiomyocyte hypertrophy (**Supplementary Figure 4**). We did not observe that deletion of *Hif1α* in resident cardiac macrophages increases resident cardiac macrophage cell death (**Supplementary Figure 5**). These findings indicate that deletion of *Hif1α* in resident cardiac macrophages is sufficient to recapitulate many of the phenotypes observed in mice lacking *Hif1α* in all monocytes and macrophages and suggest that hypoxia sensing in cardiac resident macrophages regulates the specification of infiltrating monocytes and subsequently impacts post-MI LV remodeling.

### Genetic lineage tracing reveals that Arg1^+^ macrophages differentiate into transcriptionally distinct macrophage subsets that persist throughout MI

Trajectory analysis predicted that Arg1^+^ macrophages are an intermediate population that gives rise to additional macrophage subsets. To explore this possibility, we bred *Arg1^tdT-CreERT^*^2^ *Rosa26^lsl-ZsGreen^* mice^35,39^ (Arg1ZsGr) to track Arg1^+^ cells and their progeny after MI. As Arg1^+^ macrophages are specified early after MI, we administered tamoxifen (60mg/kg IP) to Arg1ZsGr mice prior to and immediately after I/R (days −1, 0, +1) to label and track the fates of Arg1^+^ macrophages. Tissues were collected 2, 7, and 30 days after I/R for IHC and FACS. ZsGr^−^ and ZsGr^+^ monocytes and macrophages were collected 2 and 30 days after I/R for scRNAseq (**Fig. 5a**). ARG1 co-localized with ZsGr^+^ cells specifically within the infarct 2 days after I/R. At later time points, no ARG1 expression was observed in ZsGr^+^ cells (**Extended Data Fig. 5a**-b). ZsGr^+^ monocytes and macrophages persisted 2, 7, and 30 days after I/R via FACS, indicating that cells derived from Arg1^+^ macrophages are maintained throughout MI (Fig. 5b). This was corroborated by IHC co-staining of CD68 with ZsGr^+^ cells within the infarct (**Fig. 5c-d**). ZsGr^+^CD68^+^ cells were also found in the remote zone 7 and 30 days after MI (**Fig. 5c-d**).

**Fig. 5.**
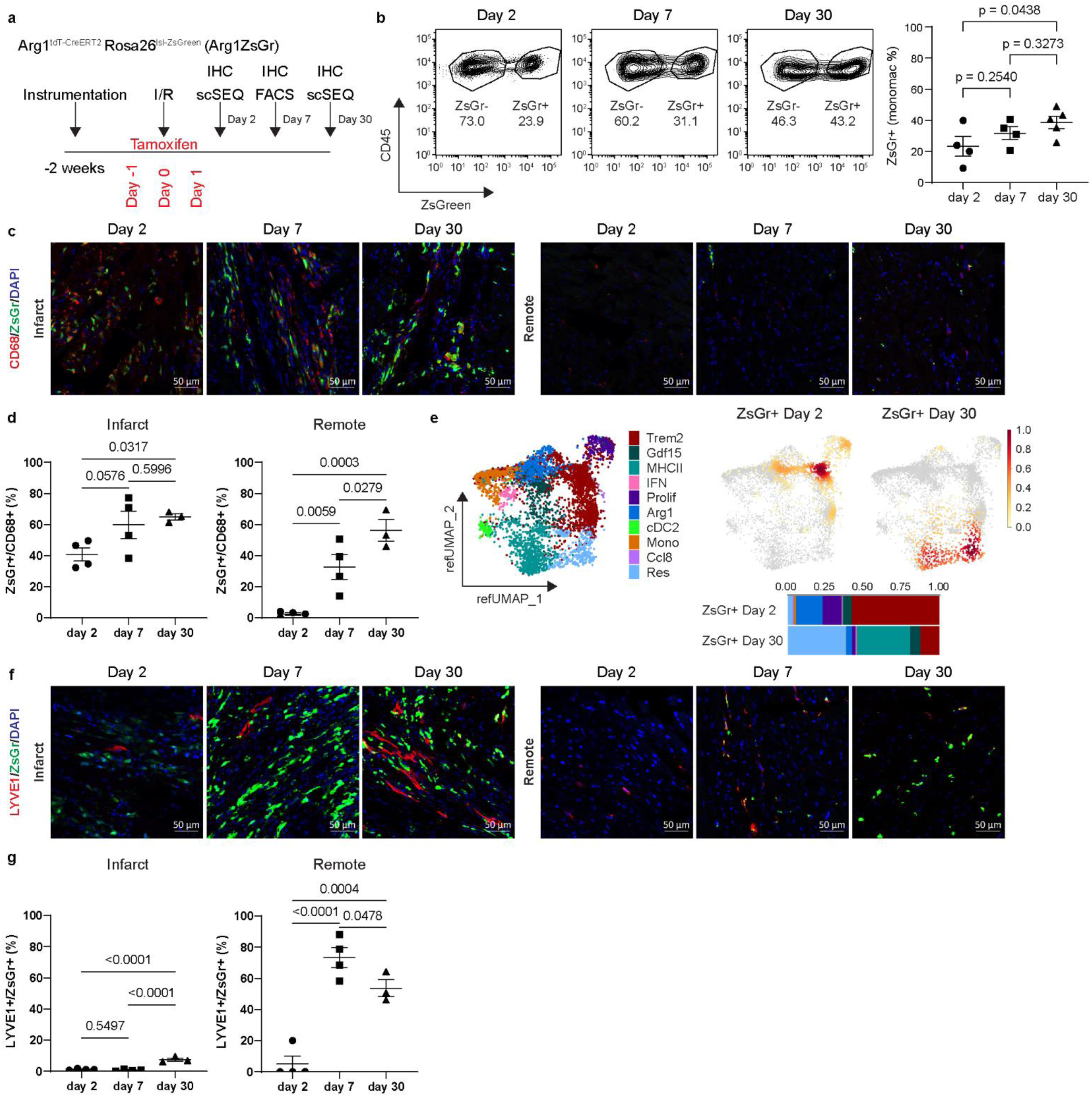
The Arg1 macrophage lineage is maintained through MI and differentiates into various macrophage subtypes. **a,** Experimental outline of the I/R mouse model of MI in Arg1ZsGr mice used to lineage trace Arg1 macrophages. Closed chest I/R was performed 2 weeks after instrumentation, and tissues collect 2, 7, and 30 days after I/R for FACS and IHC. ZsGr- and Zsgr+ monocytes and macrophages were collected 2 and 30 days after I/R for scSEQ. 60 mg/kg tamoxifen was IP injected −1, 0, and +1 days relative to the time of injury. **b,** Representative FACS and quantification of ZsGr^−^ and ZsGr^+^ macrophages in Arg1ZsGr mice 2, 7, and 30 days after I/R. A monocyte/macrophage back gate was used. **c,** Representative 20x confocal images of Arg1 macrophage lineage IHC in the infarct and remote zones of Arg1ZsGr mice 2, 7, and 30 days after I/R. CD68 staining is in red, ZsGr is in green, and DAPI is in blue. **d,** Quantification of IHC in c. Data is displayed as the percentage of ZsGr^+^CD68^+^ cells out of total CD68^+^ cells per 20x field. **e,** Annotated reference UMAP, gaussian kernel density estimate plots, and stacked bar graph of ZsGr^+^ scSEQ data 2 and 30 days after I/R in Arg1ZsGr mice. Data was mapped to a reference MI data set of monocytes, macrophages, and dendritic-like cells 3, 7, 13, and 28 days after I/R. **f,** Representative 20x confocal images of ZsGr^+^ resident macrophages in the infarct and remote zones of Arg1ZsGr mice 2, 7, and 30 days after I/R. Resident macrophage marker LYVE1 is in red, ZsGr is in green, DAPI is in blue. **g,** Quantification of IHC in f. Data is displayed as the percentage of ZsGr+ cells that are LYVE1+Zsgr+ per 20x field. P-values are determined using the ordinary one-way ANOVA for multiple comparisons.

scRNAseq of ZsGr^−^ and ZsGr^+^ monocytes and macrophages was then performed and the data mapped to our reference MI data set to maintain consistent annotations across experiments (**Extended Data Fig. 5c**-i). Arg1 expression was highest in ZsGr^+^ cells at day 2, consistent with the premise that ZsGr labels Arg1^+^ macrophages and can be used to track their progeny (**Extended Data Fig. 5e**). Composition analysis showed that at day 2 after MI ZsGr^+^ cells consisted primarily of Arg1^+^, proliferating, Trem2^+^, and Gdf15^+^ macrophages. At day 30 after MI, ZsGr^+^ cells were comprised of Trem2^+^ macrophages, Gdf15^+^, MHC-II^hi^, and resident-like macrophages (**Fig. 5e**). The contribution of Arg1^+^ macrophages to a resident-like population was validated by IHC for LYVE1 (resident macrophage marker). LYVE1^+^ZsGr^+^ cells were detected in the infarct 30 days after MI and in the remotes zone 7 and 30 days after MI (**Fig. 5f-g**). These findings indicate that Arg1^+^ macrophages specified early after MI differentiate into multiple downstream macrophage subsets including a population that resembles cardiac resident macrophages.

### Arg1 lineage tracing using a syngeneic heart transplantation model identifies a distinct monocyte differentiation trajectory

The finding that ZsGr^+^CD68^+^ cells were present in the infarct and remote zones after MI is consistent with the premise that Arg1^+^ macrophages and their progeny constitute a previously unappreciated trajectory of macrophage differentiation. Alternatively, this observation could also be explained by transient expression of *Arg1* in cardiac resident macrophages following MI. To examine the latter possibility, we transplanted Arg1ZsGr donor hearts into syngeneic recipients and administered tamoxifen (60mg/kg IP) prior to and immediately after transplant (days −1, 0, +1) to label and track donor resident macrophages that might express *Arg1*. Consistent with the conclusion that cardiac resident macrophages do not express *Arg1* in response to I/R, we detected very few CD68^+^ZsGr^+^ cells 7 days following transplantation within the myocardium. It is important to note that uninjured hearts did contain CD68^+^ZsGr^+^ cells 7 days after tamoxifen administration (**Extended Data** Fig.6).

To completely avoid any possible labeling of cardiac resident macrophages, we repeated our Arg1^+^ lineage tracing studies using the syngeneic heart transplant model of ischemic heart injury^40–42^. Donor C57BL6 hearts were transplanted into Arg1ZsGr recipients to ensure that ZsGr^+^ cells in the donor heart would be derived from monocytes. The syngeneic heart transplant model also provides the opportunity to investigate the impact of *Hif1α* deletion in resident macrophages on the differentiation kinetics of Arg1^+^ macrophages by transplanting either control or resHif1aKO donor hearts into Arg1ZsGr recipients. Tamoxifen was administered to control and resHif1aKO donor mice (60 mg/kg gavage for 5 days) to delete *Hif1α* in donor (cardiac resident) macrophages. Donor hearts were then placed on ice for 60 minutes (ischemia) and transplanted into recipient Arg1ZsGr mice (Control>ZsGr and Hif1a>ZsGr). Tamoxifen (60mg/kg) was administered prior to and immediately after transplant (days −1, 0, +1) to label and track recipient monocyte-derived Arg1^+^ macrophages. Donor hearts were collected 5 days after transplant for IHC, FACS, and scRNAseq (**Fig. 6a**). Consistent with the existence of a lineage derived from Arg1^+^ macrophage, FACS revealed that control and resHif1aKO transplanted hearts contained both ZsGr^−^ and ZsGr^+^ macrophages (**Fig. 6b-c, Extended Data Fig. 7a**).

**Fig. 6.**
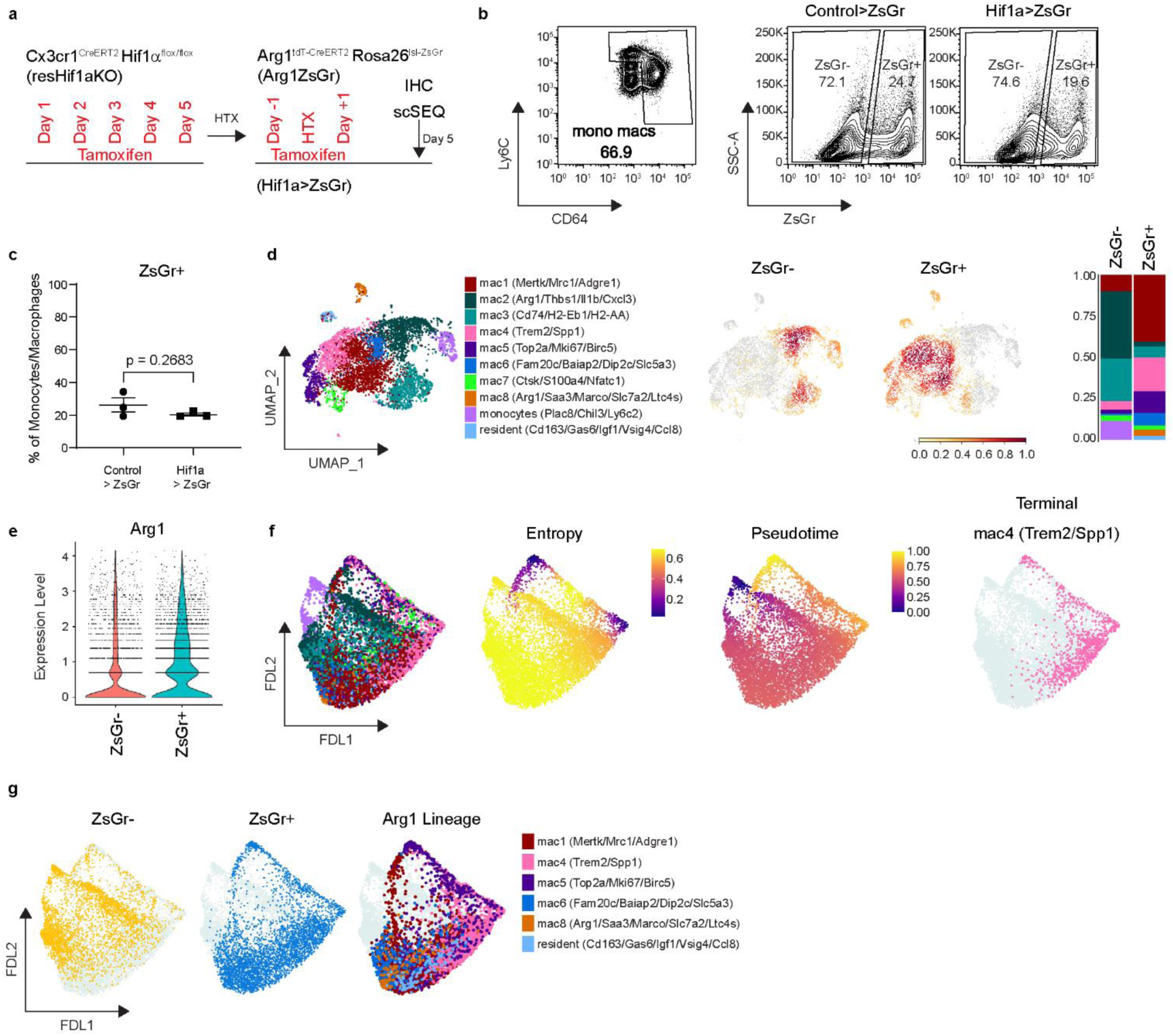
The Arg1 macrophage lineage represents a distinct monocyte to macrophage lineage during ischemic heart injury. **a,** Experimental outline of the syngeneic heart transplant model used to lineage trace Arg1^+^ macrophages when the donor heart is either a control or resident macrophage Hif1α knockout (resHif1aKO) heart. 60 mg/kg of tamoxifen was administered via gavage for 5 days in Control and resHif1aKO mice and then transplanted into Arg1ZsGr mice (Control>ZsGr or Hif1a>ZsGr). 60 mg/kg of tamoxifen was administered via gavage to recipient Arg1ZsGr mice −1, 0, and +1 days relative to the time of transplant. Tissues were collected 5 days after transplant for IHC, FACS, and scSEQ. ZsGr^−^ and Zsgr^+^ monocytes and macrophages were collected for scSEQ. **b,** Representative FACS of ZsGr^−^ and Zsgr^+^ monocytes and macrophages in Control>ZsGr and Hif1a>ZsGr mice. **c,** Quantification of the percentage of ZsGr^+^ monocytes and macrophages in Control>ZsGr and Hif1a>ZsGr mice. **d,** Annotated UMAP, gaussian kernel density estimate plots, and stacked bar graph of scSEQ data in Control>ZsGr and Hif1a>ZsGr mice, split by ZsGr^−^ and ZsGr^+^ cells. **e,** Violin plot of *Arg1* expression in Control>ZsGr and Hif1a>ZsGr mice, split by ZsGr^−^ and ZsGr^+^ cells. **f,** FDL representation of Palantir analysis displaying all the cells as annotated in d, entropy and pseudotime scores, and the predicted terminal state mac4 (Trem2/Spp1). **g,** FDL layout of Palantir analysis split by ZsGr^−^ and ZsGr^+^ cells, and cells overrepresented in ZsGr^+^ libraries (Arg1 Lineage; mac1, mac4, mac5, mac6, mac8, and resident macrophages.) P-values are determined using a two tailed t-test assuming equal variance.

We then generated scRNAseq libraries from ZsGr^−^ and ZsGr^+^ monocytes and macrophages in Control>ZsGr and Hif1a>ZsGr mice. Clustering identified 10 transcriptionally distinct subsets of monocytes and macrophages (**Extended Data Fig. 7b**-f). We first compared ZsGr^−^ to ZsGr^+^ cells within this dataset. Composition analysis revealed that ZsGr^+^ cells were transcriptionally distinct from ZsGr^−^ cells. ZsGr^+^ cells displayed a greater abundance of the following subsets: mac1 (*Mertk*, *Mrc1*, *Adgre1*), mac4 (*Trem2*, *Spp1*), mac5 (*Top2a*, *Mki67*, *Birc5*), mac6 (*Fam20c*, *Baiap2*, *Dip2c*, *Slc5a3*), mac8 (*Arg1*, *Saa3*, *Marco*, *Slc6a2*, *Ltc4s*), and resident-like macrophages (*Cd163*, *Gas6*, *Igf1*, *Vsig4*, *Ccl8*). These data indicate that macrophages differentiating through the Arg1 lineage differ from other monocyte-derived macrophages (**Fig. 6d**). These data also recapitulate several key findings observed in our Arg1 lineage tracing studies performed in the MI model. Specifically, the presence of Trem2^+^, proliferating, and resident-like macrophages. Consistent with the expectation that ZsGr would label Arg1^+^ macrophages and their progeny, we also observed higher *Arg1* expression in ZsGr^+^ cells (**Fig. 6e**).

Palantir trajectory analysis^32^ was performed on ZsGr^−^ and ZsGr^+^ cells, with monocytes being selected as the starting cell type (**Extended Data Fig. 8a**-d). This analysis suggested that the Arg1^+^ macrophages are an intermediate macrophage state and that the Arg1^+^ macrophage lineage diverged from other monocytes and macrophages present within the transplanted heart. This analysis also predicted that mac4 (*Trem2*, *Spp1*) represented a terminal differentiated cell state (**Fig. 6f-g**). Collectively, these findings indicate that Arg1^+^ macrophages are derived from monocytes that infiltrate the heart after ischemic injury and subsequently differentiate into several different subsets including Trem2^+^, proliferating, and resident-like macrophages.

### *Hif1α* in resident cardiac macrophages governs progression through the Arg1^+^ macrophage lineage

Our studies have shown that *Hif1α* deletion in resident macrophages increases the abundance monocyte-derived Arg1^+^ macrophages. To ascertain the mechanistic basis by which Arg1^+^ macrophages accumulate in this setting, we leveraged our scRNAseq data from the above syngeneic heart transplant model to investigate progression through the Arg1^+^ macrophage lineage. We compared ZsGr^+^ macrophages from control and resHif1aKO donor hearts that were transplanted into Arg1ZsGr mice (Control>ZsGr and Hif1a>ZsGr) (**Fig. 6a**). FACS and IHC analysis of ZsGr^+^ monocytes and macrophages revealed that Hif1a>ZsGr had a higher amount TDT^+^ cells compared to control 5 days after transplant (**Fig. 7a-b, Extended Data Fig. 7a**). These data indicate increased abundance of ZsGr^+^ cells expressing *Arg1* in resHif1aKO donor hearts compared to control donor hearts consistent with our MI data showing accumulation of Arg1^+^ macrophages when *Hif1α* is deleted from resident macrophages.

**Fig. 7.**
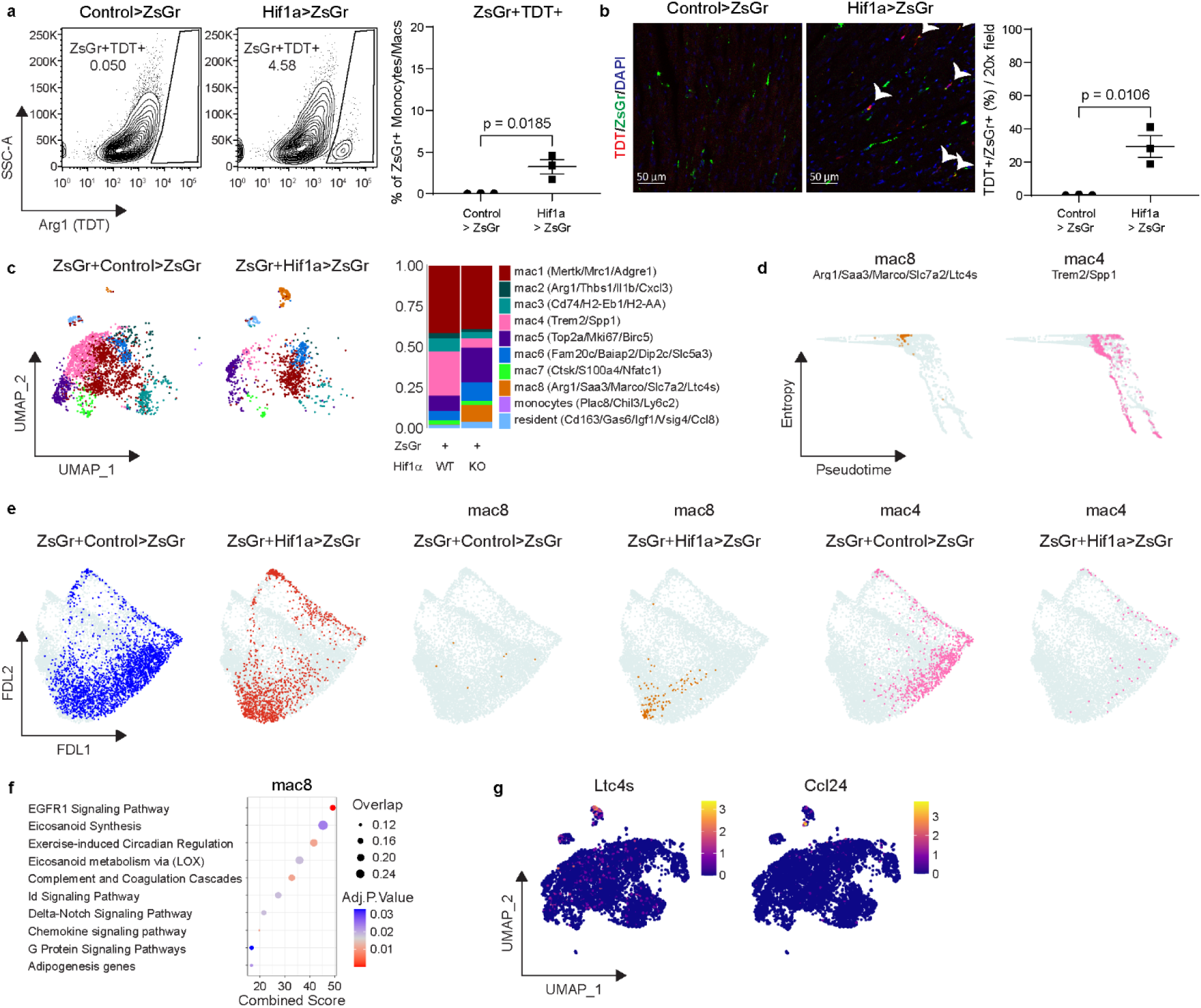
*Hif1α* activity in resident cardiac macrophages regulates progression through the Arg1 lineage. **a,** Representative FACS of ZsGr^+^tdT^+^ monocytes macrophages in Control>ZsGr and Hif1a>ZsGr mice. ZsGr^+^ monocytes and macrophages were used as a back gate. Quantification is displayed as the percentage of ZsGr^+^ monocytes and macrophages that are TDT^+^. **b,** Representative 20x confocal images of endogenous ZsGr (green) and TDT (red) IHC in Control>ZsGr and Hif1a>ZsGr mice. DAPI is in blue. Co-staining of ZsGr with TDT is highlighted by white arrows. Quantification is displayed as the percentage of ZsGr^+^ cells that are ZsGr^+^TDT^+^. **c,** Annotated UMAP and stacked bar graph of ZsGr^+^ cells split by Control>ZsGr and Hif1a>Zsgr conditions. **d,** Plots of entropy vs pseudotime scores from Palantir analysis for Arg1^+^ (mac8) and Trem2^+^ (mac4) macrophages. **e,** FDL layout of Palantir analysis of ZsGr^+^ cells split by Control>ZsGr (blue) and Hif1a>ZsGr (red) conditions. Populations of interest (mac8, orange, and mac4, pink) plotted on a FDL layout split by Control>ZsGr and Hif1a>ZsGr conditions. **f,** Dot plot of the top 10 upregulated pathways (Enrichr Wikipathways 2019 Mouse) in mac8(Arg1/Saa3/Marco/Slc7a2/Ltc4s) cells ordered by the combined score. **g,** Gene expression of Ltc4s (Eicosanoid Synthesis term) and Ccl24 (Chemokine signaling pathway term) plotted on a UMAP. P-values are determined using a two tailed t-test assuming equal variance.

To determine whether progression through the Arg1^+^ macrophage lineage is interrupted in resHif1aKO donor hearts, we next compared the composition of ZsGr^+^ cells between control and resHif1aKO donor hearts. resHif1aKO donor hearts displayed markedly lower proportions of Trem2^+^ macrophages (mac 4) and greater proportions of proliferating (mac5) and Arg1^+^ macrophages (mac8) compared to control donor hearts (**Fig. 7c**). Palantir trajectory analysis predicted that Arg1^+^ macrophages (mac8) represent an intermediate state (high entropy, low pseudotime) and Trem2^+^ macrophages (mac 4) represent a terminal state (low entropy, high pseudotime) (**Fig. 7d**). Visualization of ZsGr^+^ cells within control and resHif1aKO donor hearts in trajectory space signaled a bias towards Trem2^+^ macrophages (terminal state) and Arg1^+^ macrophages (intermediate state), respectively (**Fig. 7e**). In addition, Arg1^+^ macrophages displayed inflammatory gene signatures such as eicosanoid synthesis and chemokine signaling (**Fig. 7f-g**). Collectively, these findings demonstrate that Hif1α signaling in resident cardiac macrophage regulates the differentiation of Arg1^+^ macrophages. *Hif1α* deletion in resident cardiac macrophages resulted in the accumulation of Arg1^+^ macrophages with inflammatory potential by impeding their differentiation.

## Discussion

Here, we provide evidence that *Hif1α* deletion in monocytes and macrophages led to accelerated LV remodeling following MI and resulted in an increase in a subset of macrophages marked by *Arg1* mRNA and protein expression located within the infarct. Arg1^+^ macrophages displayed an inflammatory gene signature, were derived from monocytes, and specified early during monocyte to macrophage differentiation in the context of ischemic injury. Mechanistically, Hif1α signaling in resident macrophages served as a cell extrinsic determinant regulating the abundance of Arg1^+^ macrophages. Lineage tracing of Arg1^+^ macrophages in mouse models of I/R showed that Arg1^+^ macrophages differentiate into downstream macrophage subsets that included Trem2^+^, Gdf15^+^, MHC-II^hi^, and resident-like macrophages. This Arg1 macrophage lineage was transcriptionally distinct from other trajectories of monocyte to macrophage differentiation. Finally, we observed that *Hif1α* deletion in resident cardiac macrophages interrupted the progression of monocytes through the Arg1^+^ macrophage lineage.

There are conflicting reports pertaining the function of Hif1α signaling in myeloid cells following cardiac injury. *Hif1α^flox/flox^ LysM^Cre^* mice subjected to transverse aortic constriction displayed increased cardiac fibrosis and a reduction in ejection fraction^43^. Conversely, a separate manuscript demonstrated that *Hif1α^flox/flox^ LysM^Cre^* mice displayed decreased infarct size and increased ejection fraction following permanent occlusion MI^44^. It is possible that differences in injury models (I/R vs coronary ligation), cellular specificity, or temporal control of *Hif1α* deletion may account for differences in phenotypes. Importantly, *LysM^Cre^* is active during development and throughout the myeloid compartment, including neutrophils^44,45^.

We found that *Hif1α* deletion in monocytes and macrophages did not influence total macrophage numbers in response to MI, consistent with a previous study^44^. However, scRNAseq revealed that *Hif1α* deletion increased the abundance of Arg1^+^ macrophages and influenced monocyte to macrophage differentiation after MI. *Hif1α* deletion in cardiac resident macrophages and not monocyte-derived macrophages mediated these phenotypes. These data indicate that aberrant hypoxia sensing in resident cardiac macrophages has a non-cell autonomous effect on monocyte specification. This finding mirrors the observation that depletion of CCR2^−^ cardiac resident macrophages prior to MI results in increased Arg1^+^ macrophages within the infarct^5^. Cardiac resident macrophage death represents an important event following myocardial infarction that may impact outcomes^14^. However, we observed no difference in cell death rates of resident cardiac macrophages in resHif1aKO mice suggesting that cardiac resident macrophages are influencing monocyte differentiation through a paracrine mechanism that remains to be determined.

Transcriptional profiling indicates that Arg1^+^ macrophages possess pro-inflammatory properties. Notably, Arg1^+^ macrophages displayed enriched expression of genes involved in eicosanoid synthesis and cytokine expression. Eicosanoids, including leukotriene C4, are pro-inflammatory, induce cardiomyocyte death, and are associated with worsened remodeling^46^. These findings contrast with prior literature arguing that *Arg1* is a marker of the M2 macrophage phenotype^47–49^. It is essential to highlight that the *in vivo* relevance of the M1/M2 paradigm is uncertain^50^. Consistent with our findings, *Arg1* expressing macrophages are deleterious and contribute to enhanced fibrosis in mouse bleomycin lung injury and MI^51,52^.

Our findings provide key new insights regarding mechanisms of monocyte-derived macrophage differentiation. Through multiple approaches we showed that Arg1^+^ macrophages are derived from monocytes, are initially specified within the coronary vascular compartment prior to extravasation into the myocardium, and differentiate into multiple downstream macrophages subsets. The transcriptional signature and lineage trajectory of Arg1^+^ macrophages was distinct from interferon-activated macrophages, demonstrating that monocytes commit to unique trajectories upon recruitment to the ischemic heart. *Hif1α* deletion in resident macrophages halted the progression through the Arg1^+^ macrophage lineage resulting in accumulation of pro-inflammatory intermediates. These insights are paradigm shifting in that they uncover that monocyte fate decisions are initiated within the vascular compartment and that transitional macrophage subsets exert unique functions and later differentiate into functionally distinct subsets of macrophages.

Our study is not without limitations and much remains to be learned. The paracrine signaling pathway by which cardiac resident macrophages influence the differentiation of Arg1^+^ macrophages remains to be determined. Furthermore, the mechanistic basis by which Arg1^+^ macrophages are specified within the vasculature is also unclear. The functional requirement of Arg1^+^ macrophages in cardiac pathology must be inferred based on data regulating progression through this lineage as it is not possible to deplete Arg1+ macrophages without impacting downstream macrophage subsets (Trem2^+^, Gdf15^+^, MHC-II^hi^, resident-like macrophages), which may have differing functions. Finally, the role of *Hif1α* deletion in monocytes and monocyte-derived macrophages in post-MI LV remodeling requires further investigation.

In conclusion, we demonstrate that hypoxia sensing in cardiac resident macrophages has a protective role following MI and modulates monocyte fate decisions. We identified unique and targetable lineages of monocyte differentiation within the heart and show that hypoxia sensing in resident cardiac macrophages is essential for progression through the Arg1^+^ lineage. Collectively, our findings provide new insights into the complexity of monocyte differentiation during ischemic heart injury.

## Acknowledgments

KJL is supported by the Washington University in St. Louis Rheumatic Diseases Research Resource-Based Center grant (NIH P30AR073752), the National Institutes of Health [R01 HL138466, R01 HL139714, R01 HL151078, R01 HL161185, R35 HL161185], Leducq Foundation Network (#20CVD02), Burroughs Wellcome Fund (1014782), and Children’s Discovery Institute of Washington University and St. Louis Children’s Hospital (CH-II-2015-462, CH-II-2017-628, PM-LI-2019-829), Foundation of Barnes-Jewish Hospital (8038-88), and generous gifts from Washington University School of Medicine. FFK was supported by the National Institutes of Health (5T32GM007067-45, 5T32GM007067-46, 5T32HL134635-05). We thank the Genome Technology Access Center at the McDonnell Genome Institute at Washington University School of Medicine for help with genomic analysis. The Center is partially supported by NCI Cancer Center Support Grant #P30 CA91842 to the Siteman Cancer Center. We are grateful to Richard Locksley (UCSF) for providing us with *Arg1^tdT-CreERT^*^2^ mice. This publication is solely the responsibility of the authors and does not necessarily represent the official view of NCRR or NIH.

## Author Contributions

FFK performed IHC staining/imaging/quantification, FACS analysis and sorting, qPCR, scRNAseq analysis, trajectory analysis, and spatial transcriptomic analysis. CJW and JMN performed closed chest ischemia reperfusion surgeries. AK performed echocardiography. KJL quantified echocardiography data. ALB performed FACS sorting and 10x library preparation for macHif1aKO mice and contributed to design and interpretation of FACS experiments. ALK optimized the *Ccr2^ERT2Cre^ Rosa26^lsl-tdT^* recombination strategy, prepared/analyzed/annotated the reference MI scRNAseq dataset, and contributed to bioinformatics analyses and IHC. JMA and LL contributed to bioinformatics analyses. WL performed cervical syngeneic heart transplant and 2-photon intravital imaging. HD performed abdominal syngeneic heart transplant surgeries. BK contributed to design and interpretation of heart transplant experiments. VP contributed to design and interpretation of Cytek FACS experiments. FFK made all figures. FFK and KJL drafted the manuscript. All authors contributed to the experimental design, analysis, and interpretation as well as manuscript production. KJL is responsible for all aspects of this manuscript including experimental design, data analysis, and manuscript production. All authors approve the final version of the manuscript.

## Methods

### Animal Models

Mus musculus were housed in ultraclean rodent barrier facility with veterinarians from the Washington University Division of Comparative Medicine on call or on site for animal care. All procedures were performed in accordance with IUCAC protocols. Strains used were *Cx3cr1^ERT2Cre^* ^19^ (JAX #020940), *Hif1α^flox/flox^* ^20^ (JAX #007561), *Rosa26^lox-stop-lox^ ^tdTomato^* ^39^ (JAX #007914), *Ccr2^ERT2Cre^* ^53^, *Arg1^YFP^* ^38^ (JAX #015857), *Arg1^tdT-CreERT^*^235^ (from the Richard M Locksley lab), *Rosa26^lox-stop-lox^ ^ZsGreen^* ^39^ (JAX #007906), on the C57BL/6 background. Experiments were performed on mice 2 months to 10 months of age. Similar numbers of male and female mice were used for experiments. For Cre recombination in all monocytes and macrophages, *Cx3cr1^ERT2Cre^ Hif1α^flox/flox^ Rosa26^lox-stop-lox^ ^tdTomato^* mice were given tamoxifen food pellets (Envigo TD.130857) for the entirety of the experiment. For Cre recombination in other lines, mice were given either IP injection or gavage of 60 mg/kg of tamoxifen (Millipore Sigma T5648) at the indicated frequency and timing.

### Closed-chest Ischemia Reperfusion Injury

Mice were anesthetized, intubated, and mechanically ventilated. The heart was exposed, and a suture placed around the proximal left coronary artery. The suture was threaded through a 1 mm piece of polyethylene tubing to serve as the arterial occlude. Each end of the suture was exteriorized through the thorax. The skin was closed, and mice were given a 2-week recovery period prior to induction of ischemia. After 2 weeks, the animals were anesthetized and placed on an Indus Mouse Surgical Monitor system to accurately record ECG during ischemia. Ischemia was induced after anesthetizing the animals. Tension was exerted on suture ends until ST-segment elevation was seen via ECG. Following 90 minutes of ischemia time, tension was released, and the skin was then closed.

### Echocardiography

Mice were sedated with Avertin (0.005 ml/g) and 2D and M-mode images were obtained in the long and short axis views 4 weeks after MI using the VisualSonics770 Echocardiography System. Left diastolic volume, left systolic volume, akinetic area, and total LV area was measured using edge detection and tracking software (VivoLab). Ejection fraction was calculated as the (left diastolic volume – left systolic volume)/ left diastolic volume. Akinetic percentage was calculated as the akinetic area / total LV area.

### Syngeneic Heart Transplant and 2-photon Intravital Imaging

For syngeneic heart transplant of *Cx3cr1^ERT2Cre^ Hif1αflox/flox* into *Arg1tdT-CreERT2 Rosa26lox-stop-lox ZsGreen* mice or *Arg1tdT-CreERT2 Rosa26lox-stop-lox ZsGreen* into wildtype in the abdominal cavity, an established syngeneic heterotopic cardiac transplant protocol was used^40–42^. Donor mice were perfused with heparinized saline through the infrarenal aorta. Cardiac grafts were harvested from donor mice and left in saline on ice for 1 hour for cold ischemia. Recipient mice were anesthetized with ketamine (100 mg/kg) and xylazine (10 mg/kg) and maintained with 1%-2% isoflurane. The recipient infrarenal aorta and inferior vena cava were connected to the donor ascending aorta and pulmonary artery, respectively.

For *Arg1^YFP^* intravital 2-photon imaging following syngeneic heart transplant, established protocols were used^8,54,55^. Cardiac grafts were harvested from donor wild type mice and transplanted into the cervical position of recipient *Arg1^YFP^*mice after 1 hour of cold ischemia. The donor ascending aorta and pulmonary artery were connected to the recipient right common carotid artery and right external jugular vein. 24 hours after syngeneic heart transplantation, mice were anesthetized with ketamine (80mg/kg) and xylazine (10 mg/kg), intubated, and connected to a mouse ventilator. 655-nm non targeted Q dots (Thermo Fisher Scientific) were injected through the intravenous route into recipient mice prior to imaging. The heart was fixed to imaging chamber and imaged for a duration of 15 minutes. Time-lapse imaging was performed with a custom-built 2-photon microscope running ImageWarp version 2.1 acquisition software (A&B Software). Imaging is terminal and mice were euthanized.

### Histology

4 weeks after MI, mice were euthanized, and hearts perfused with PBS. Hearts were fixed overnight with 4% PFA in PBS at 4⁰C, cut into thirds on the transverse plane, placed into histology cassettes for paraffin embedding, and dehydrated in 70% ethanol in water. Hearts were paraffin embedded and cut into 4 µm sections and mounted on positively charged slides. Tissues were stained by Gomori’s Trichrome staining (Richard-Allan Scientific 87020) and WGA staining (Vector Laboratories RL-1022). Trichrome staining was imaged using a Meyer PathScan Enabler IV. Measurement of infarct length and LV length was performed in FIJI, with data quantification normalizing the length of the infarct to the length of the LV^56^. WGA staining was imaged using the 20x objective of a Zeiss LSM 700 confocal microscope. 4-5 20x fields of the border zone of the infarct were taken per mouse. 12 cross sectional (not longitudinal) cardiomyocytes per 20x field were selected at random. Measurement of cross-sectional cardiomyocyte area was performed in FIJI. Data is displayed as the average cardiomyocyte cross sectional area per mouse.

4 days after MI, mice were euthanized, and hearts perfused with PBS. Hearts were incubated in 2% 2, 3, 5-triphenyltetrazolium chloride (Millipore Sigma T8877) in PBS for 30 minutes at 37⁰C and then washed in 4% PFA in PBS for 24 hours at 4⁰C. Pictures were taken with a ZEISS SteREO Discovery.V12 Stereo Microscope. Quantification of total infarct area normalized to total heart area was performed with FIJI.

### Immunofluorescence

Mice were euthanized, and hearts perfused with PBS. Hearts were cut midway on the transverse plane and fixed overnight with 4% PFA in PBS at 4⁰C. They were then dehydrated in 30% sucrose in PBS overnight at 4⁰C. Hearts were embedded in O.C.T. (Sakura 4583) and frozen at −80⁰C for 30 minutes. Hearts were then sectioned at 15 µm using a Leica Cryostat, mounted on positively charged slides, and stored at −80⁰C. For staining, the following steps were performed at room temperature and protected from light. Slides were brought to room temperature for 5 minutes and washed in PBS for 5 minutes. Sections were then permeabilized in 0.25% Triton X in PBS for 5 minutes. Sections were then blocked in 5% BSA in PBS for 1 hour. Sections were then stained with primary antibody diluted in 1% BSA in PBS for 1 hour (rat anti CD34 abcam ab8158 1:200 dilution, rat anti CD68 BioLegend 137002 1:400 dilution, rat anti Ly6g BD Pharmingen 551459 1:100 dilution, rabbit anti Arg1 Cell Signaling Technology 93668 1:100 dilution, mouse anti a-SMA conjugated to 488 Cell Signaling Technologies 46469S 1:100 dilution, rabbit anti Lyve1 abcam ab14917 1:200 dilution, rabbit anti Vsig4 abcam ab252933 1:100 dilution). Sections were then washed 3 times 5 minutes each with PBS-Tween. After washing, the appropriate secondary antibodies were added diluted in 1% BSA in PBS for 1 hour at a 1:1000 dilution (Goat anti Rat 488 Thermo Fisher Scientific a11006, Goat anti Rat 647 Thermo Fisher Scientific a21247, Donkey anti Rabbit 647 Abcam ab150075, Chicken anti Rabbit 647 Thermo Fisher Scientific a21443, Goat anti Rabbit 555 Thermo Fisher Scientific a21428, Donkey anti Rat 488 Thermo Fisher Scientific a21208). Slides were again washed 3 times 5 minutes each with PBS-T, and then mounted with DAPI mounting media (Millipore Sigma F6057) and cover slipped. Slides were stored at 4⁰C protected from light and imaged within 1 week. For immunofluorescence staining of paraffin embedded tissues collected 4 weeks after MI, the OPAL 4-Color Manual IHC Kit (Akoya Biosciences NEL810001KT) was used. Primary antibodies used were: Anti CD34 (abcam ab8158 1:200 dilution), and anti a-SMA (Invitrogen 14-9760-82 1:500 dilution).

Slides were imaged using the 20x objective of a Zeiss LSM 700 confocal microscope. Visualization of tdTomato and ZsGreen was from the endogenous signal. For quantification, 4-5 20x fields were taken per mouse of either the remote zone, border zone, infarct zone, or myocardium as indicated. Quantification of cell numbers, area percentages, or a-SMA+ vessels were performed in FIJI or Zeiss Zen, and data is displayed as the average of each 20x field per mouse.

### Flow Cytometry

Hearts were perfused with PBS, weighed, minced, and digested for 45 minutes at 37⁰C in DMEM (Gibco 11965-084) containing 4500 U/ml Collagenase IV (Millipore Sigma C5138), 2400 U/ml Hyaluronidase I (Millipore Sigma H3506), and 6000 U/ml DNAse I (Millipore Sigma D4527). Enzymes were deactivated with HBSS containing 2% FBS and 0.2% BSA and filtered through 40-micron strainers. Cells were incubated with ACK lysis buffer (Gibco A10492-01) for 5 minutes at room temperature. Cells were washed with DMEM and resuspended in 100 µl of FACS buffer (PBS containing 2% FBS and 2mM EDTA).

Approximately 100 μl of blood was collected by cheek bleed into a tube containing 20 μl of EDTA (Corning 46-034-CI). Cells were incubated with ACK lysis buffer for 15 minutes at room temperature, washed, and ACK lysed a second time. Cells were then washed with DMEM and resuspended in 100 μl of FACS buffer.

Cells were incubated in a 1:200 dilution of fluorescence conjugated monoclonal antibodies for 30 minutes at 4⁰C. Samples were washed in FACS buffer and resuspended in a final volume of 250 µl FACS buffer. Flow cytometry and sorting was performed using a BD FACS melody.

For FACS analysis of monocytes, neutrophils, and macrophages in macHif1aKO mice, antibodies used were: CD45 PerCP/Cy5.5 (BioLegend 103132), CD64 PE/Cy7 (BioLegend 139313), Ly6g FITC (BioLegend 127606), and Ly6c APC (BioLegend 128016).

For sorting of monocytes and macrophages in macHif1aKO mice for single cell RNA sequencing, antibodies used were: CD45 PerCP/Cy5.5 (BioLegend 103132), Ly6g FITC (BioLegend 127606), Ly6c APC (BioLegend 128016), DAPI (BD Pharmingen 564907), and CD64 PE/Cy7 (BioLegend 139313).

For sorting and FACS analysis of monocytes and macrophages in *Arg1^tdT-CreERT^*^2^, Arg1-YFP, Arg1ZsGr, Control>ZsGr, and Hif1a>ZsGr mice, antibodies used were: CD45 PerCP/Cy5.5(BioLegend 103132), CD11b BV421 (BioLegend 101235), Ly6g APC/Cy7 (BioLegend 127623), Ly6c BV510 (BioLegend 128033), CD64 PE/Cy7 (BioLegend 139313).

For intravascular staining, 200 μl of CD45 APC (BioLegend 103112) was injected intravascularly at a concentration of 5μg/ml in PBS 5 minutes prior to tissue collection. Hearts were then perfused with 10 ml of PBS.

For intracellular staining of Arg1-YFP, the BioLegend Cyto-Fast Fix/Perm Buffer Set (BioLegend 426803) was used following staining for surface markers, and intracellular staining with GFP-PE (BioLegend 338003) was performed.

For assessment of resident cardiac macrophage death in resHif1aKO mice 1 day after I/R, a Cytek Aurora was used. Antibodies used were MHCII APC/Cy7 (BioLegend 107628), CCR2 BV421 (BioLegend 150605), CD64 APC (BioLegend 139306), Zombie Aqua (BioLegend 77143), Ly6c PE/Cy7 (BioLegend 128017), Ly6g FITC (BioLegend 127606), CD45 Percp/Cy5.5 (BioLegend 103132), and CD11b BV785 (BioLegend 101243).

### Single Cell RNA Sequencing Analysis

Monocytes and macrophages from macHif1a KO and control mice were sorted 5 days after I/R. Single cell RNA sequencing libraries were generated using the 10X Genomics 5’v1 platform. Libraries were sequenced using a NovaSeq at the McDonnel Genome Institute (MGI). Sequencing reads were aligned to the GRcm38 transcriptome using CellRanger (v3.1.0) from 10X Genomics.

ZsGr- and ZsGr+ monocytes and macrophages were sorted from Arg1ZsGr mice 2 and 30 days after I/R. Single cell RNA sequencing libraries were generated using the 10X Genomics 5’v1 platform. Sequencing reads were aligned to the mm10-2020-A transcriptome using CellRanger (v7.1.0)

ZsGr- and ZsGr+ monocytes and macrophages were sorted from Control>ZsGr and Hif1a>ZsGr mice 5 days after syngeneic heart transplant. Libraries were generated using the 10x 3’v3.1 platform. Sequencing reads were aligned to the mm10-2020-A transcriptome using CellRanger (v7.1.0).

Downstream data processing was performed in the R package Seurat (v4)^57^. SCtransform was used to scale and normalize the data for RNA counts. Quality control filters of genes/cell, mitochondrial reads/cell, and read counts/cell were applied as indicated (Extended Data Fig. 2b, Extended Data Fig. 5d, Extended Data Fig. 7b). PCA was used for dimensionality reduction, data was clustered at multiple resolutions, and UMAP projections were generated for data visualization. Harmony integration was used for Arg1ZsGr I/R data to account for batch effect from different time points^58^. Contaminating populations of neutrophils and non-myeloid cells were excluded from analysis. Annotation of cell populations in I/R data sets was achieved by using label transfer in Seurat, mapping the data to a reference data set of myeloid cells present in the mouse heart 3, 7, 13, and 28 days after MI^24,25^. Differential gene expression of upregulated genes in each subpopulation was generated in Seurat using FindAllMarkers with a minimum fraction (min.pct) of 0.1 and a logFC threshold of 0.1. Differential gene expression between macHif1aKO and control mice was performed through Seurat using DEseq2^59^. The top 30 genes upregulated in macHif1aKO was used to plot a Z-score of differentially expressed genes on the UMAP. Genes with a log2FC > 0.1 and adjusted p value of < 0.05 from FindAllMarkers were used as input for Wikipathways 2019 Mouse analysis and ChEA 2016 transcription factor analysis in EnrichR^60–64^. Gaussian kernel density estimation plots were generated using the Python package Scanpy (v1.8.1)^65^.

### Palantir Trajectory Analysis

The Palantir Python package was used for trajectory analysis (v1.0.0 to v1.1)^32^. Prior to inputting data to Palantir, cell cycle regression was performed using Seurat. Monocytes were used as the early and starting cell type for Palantir. Highly variable gene analysis was not used. The eigenvectors used was determined by the eigengap. The number of waypoints used was 500. The option to use the early cell as the start cell was used. The output pseudotime/entropy values, terminal state probabilities, and FDL coordinates for each cell was used to generate plots in R.

### Spatial transcriptomics

Spatial transcriptomics data of Arg1 macrophage localization was generated using a published data set of mouse MI 1, 3, 5, and 7 days after injury (GSE165857)^34^. Spatial data was analyzed using Seurat. SCTransform was used to scale and normalize RNA counts. RNA and calculated Z-scores were plotted using the function “SpatialFeaturePlot”. *Myh6*, *Tnni3*, and *Tnnt2*, were used as a cardiomyocyte gene expression profile to calculate a Z score. Spots that had less than −0.25 cardiomyocyte Z score were considered infarct, and all other spots were considered non-infarct.

### Quantitative Polymerase Chain Reaction

For assessment of recombination efficiency of the *Hif1α^flox/flox^* allele, 100,000 monocytes and macrophages from control and macHif1aKO mice were sorted into RLT buffer 5 days after closed-chest I/R. RNA was isolated using a RNeasy Micro Kit (Qiagen 74004). cDNA was generated using the iScript Reverse Transcription Supermix (Bio-Rad 1708841) kit. qPCR reactions were prepared with PowerUP SYBR Green Master Mix (Thermo Fisher Scientific A25742). Reference control *36b4* primers used were: Forward: ATCCCTGACGCACCGCCGTGA, Reverse: TGCATCTGCTTGGAGCCCACGTT. *Hif1α* primers used were: Forward: TTCTGTTATGAGGCTCACCATC, Reverse: TCTGTGCCTTCATCTCATCTTC. Primers were purchased from IDT. qPCR was performed using a QuantStudio 3 (Thermo Fisher Scientific). *Hif1α* expression was normalized to *36b4* expression.

### Statistical Analysis

For analysis of echocardiography, histology, immunofluorescence, and FACS, two tailed t-test assuming equal variance was performed in Excel or Prism. For statistical comparison between more than 2 conditions, the ordinary one-way ANOVA for multiple comparisons was performed in Prism. P < 0.05 was a priori considered statistically significant. Data was plotted using Prism software.

## Data Availability

Single Cell RNA sequencing data will be available on the Gene Expression Omnibus at the time of publication.

## Code Availability

R and Python scripts are available on reasonable request.

---

**Extended Data Fig. 1.**
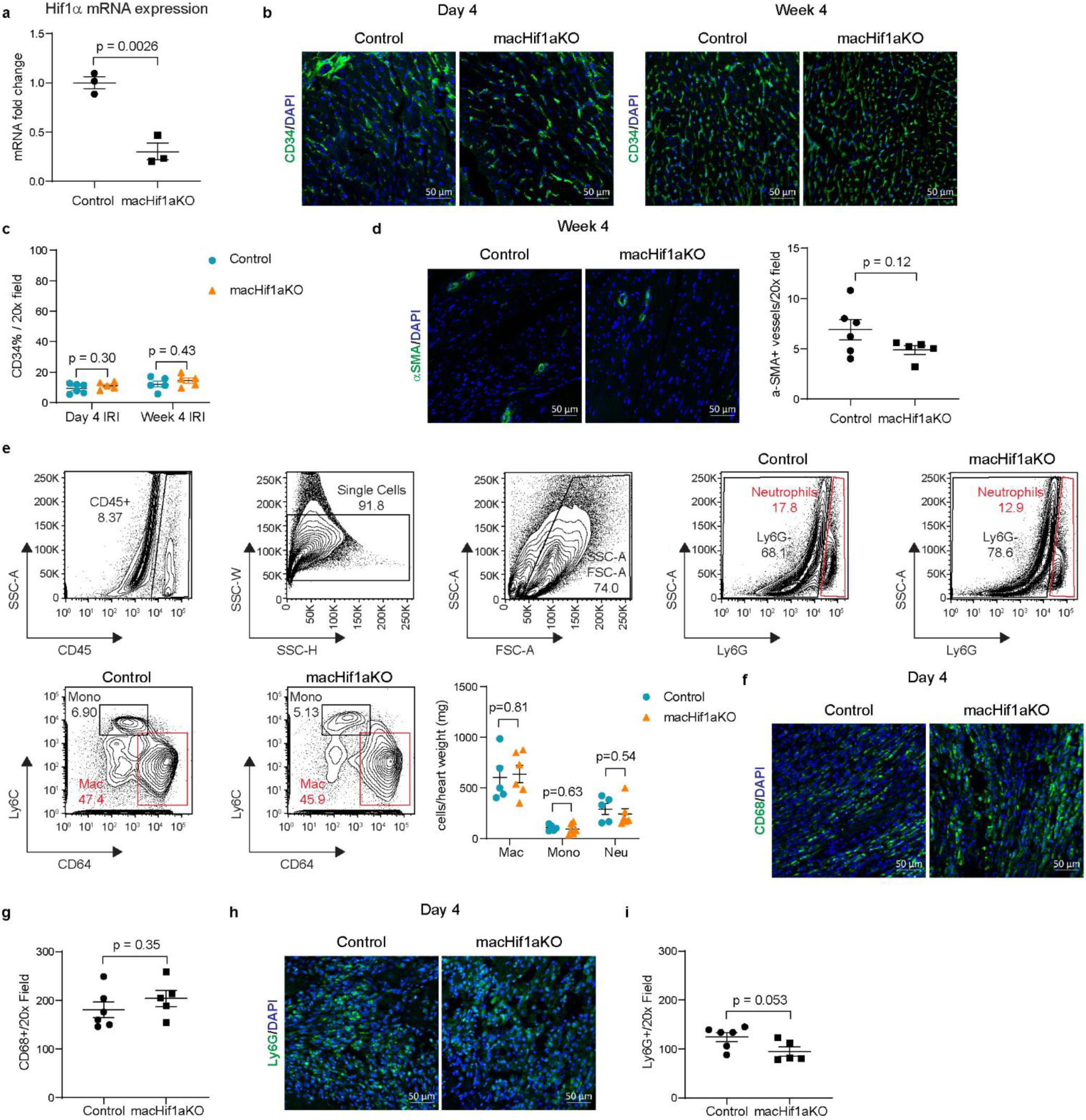
Hif1α deletion in monocytes and macrophages does not impact angiogenesis or the myeloid compartment after MI. **a,** qPCR of *Hif1α* mRNA expression in sorted monocytes and macrophages from Control and macHif1aKO mice 5 days after I/R. **b,** Representative 20x confocal images of CD34 (green) IHC staining within the border zone of Control and macHif1aKO mice 4 days and 4 weeks after I/R. DAPI is in blue. **c,** Quantification of b, displayed as the percentage of CD34 signal per 20x field. **d,** Representative 20x confocal images of αSMA (green) IHC in the border zone of Control and macHif1aKO mice 4 weeks after I/R. DAPI is in blue. Quantification is displayed as the total number of αSMA^+^ vessels per 20x field. **e,** FACS from Control and macHif1aKO mice 5 days after I/R. Quantification shows cell numbers normalized to mg of heart tissue used for FACS. **f,** Representative 20x confocal images of macrophage IHC within the infarct of Control and macHif1aKO mice 4 days after I/R. CD68 staining is in green, DAPI is in blue. **g,** Quantification of f displayed as the total number of CD68^+^ cells per 20x field. **h,** Representative 20x confocal images of neutrophil IHC within the infarct of Control and macHif1aKO mice 4 days after I/R. Ly6G staining is in green and DAPI staining is in blue. **i,** Quantification of h displayed as the total number of Ly6G^+^ cells per 20x field. P-values are determined using a two tailed t-test assuming equal variance.

**Extended Data Fig. 2.**
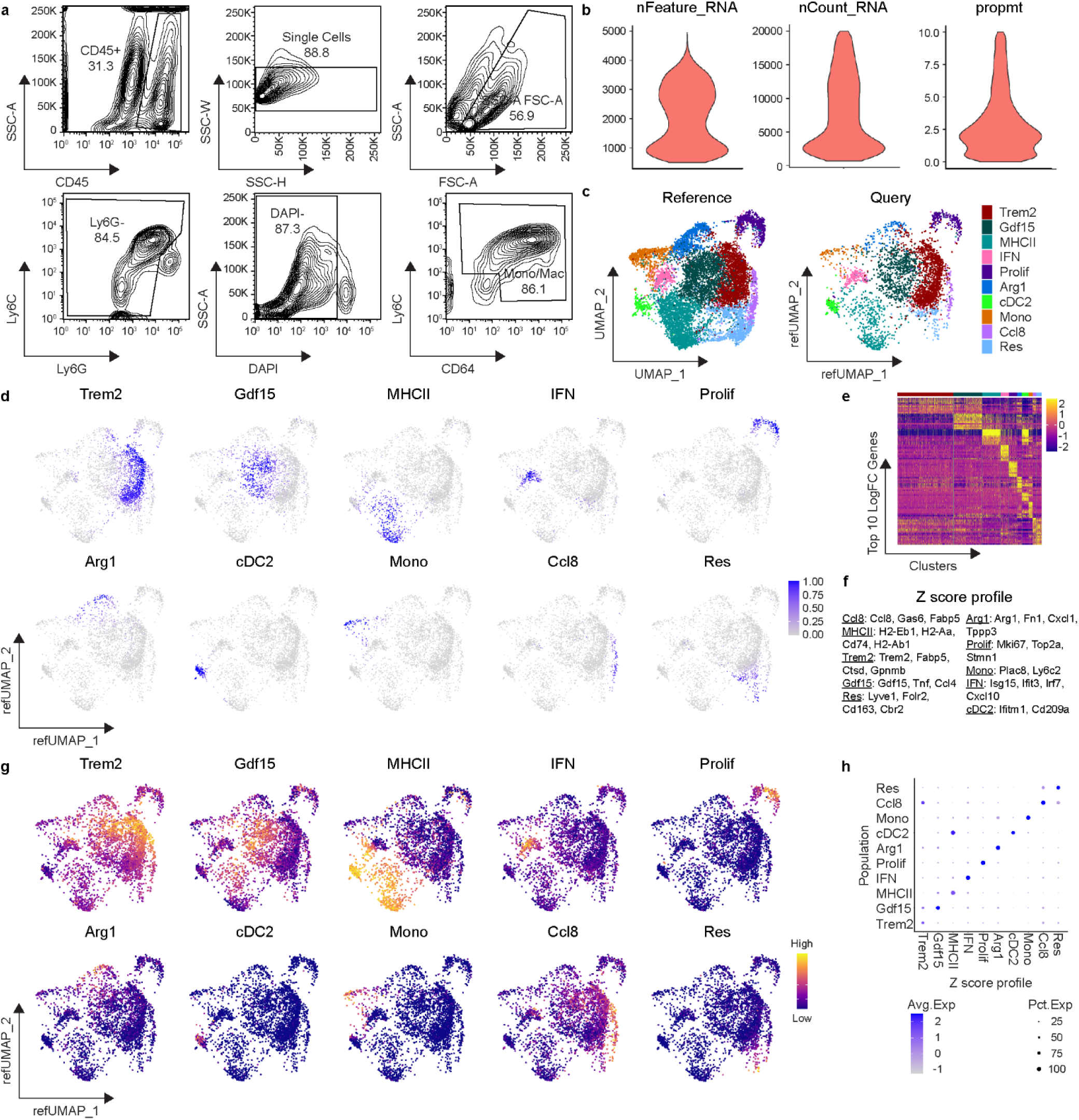
Single cell RNA sequencing and reference mapping of cardiac monocytes, macrophages, and dendritic-like cells 5 days after MI. **a,** FACS gating strategy to isolate monocytes, macrophages, and dendritic-like cells from Control and macHif1aKO mice 5 days after I/R for use in scSEQ. Cells were gated for leukocytes (SSC-A vs CD45), single cells (SSC-W vs SSC-H), debris exclusion (SSC-A vs FSC-A), neutrophil exclusion (Ly6C vs Ly6G), live cells (SSC-A vs DAPI) and monocytes/macrophages (Ly6C vs CD64). **b,** Violin plots representing quality control of scSEQ data post quality control. Cells with greater than 500 and less than 5000 genes (nFeature_RNA), read counts less than 20,000 (nCount_RNA), and proportion of transcripts mapping to mitochondrial genes less than 10 percent (prompt) were used for downstream analysis. **c,** UMAP projection and annotations from the reference MI scSEQ data set (monocytes, macrophages, and dendritic-like cells 3, 7, 13, and 28 days after I/R) and the query dataset (Control and macHif1aKO monocytes, macrophages, and dendritic-like cells 5 days after I/R) after reference mapping in Seurat. Query dataset is plotted on a reference UMAP (refUMAP) **d,** Mapping scores of each subpopulation projected onto refUMAP of the query data set (Control and macHif1aKO). **e,** Heatmap of the top 10 upregulated genes in each subpopulation across the entire Control and macHif1aKO query data set. **f,** Z-score profile genes of each subpopulation from the reference data set. **g,** Z-scores from f projected onto refUMAP of the query data set (Control and macHif1aKO. **h,** Z-scores from f compared to each subpopulation in the query data set represented as a dot-plot.

**Extended Data Fig. 3.**
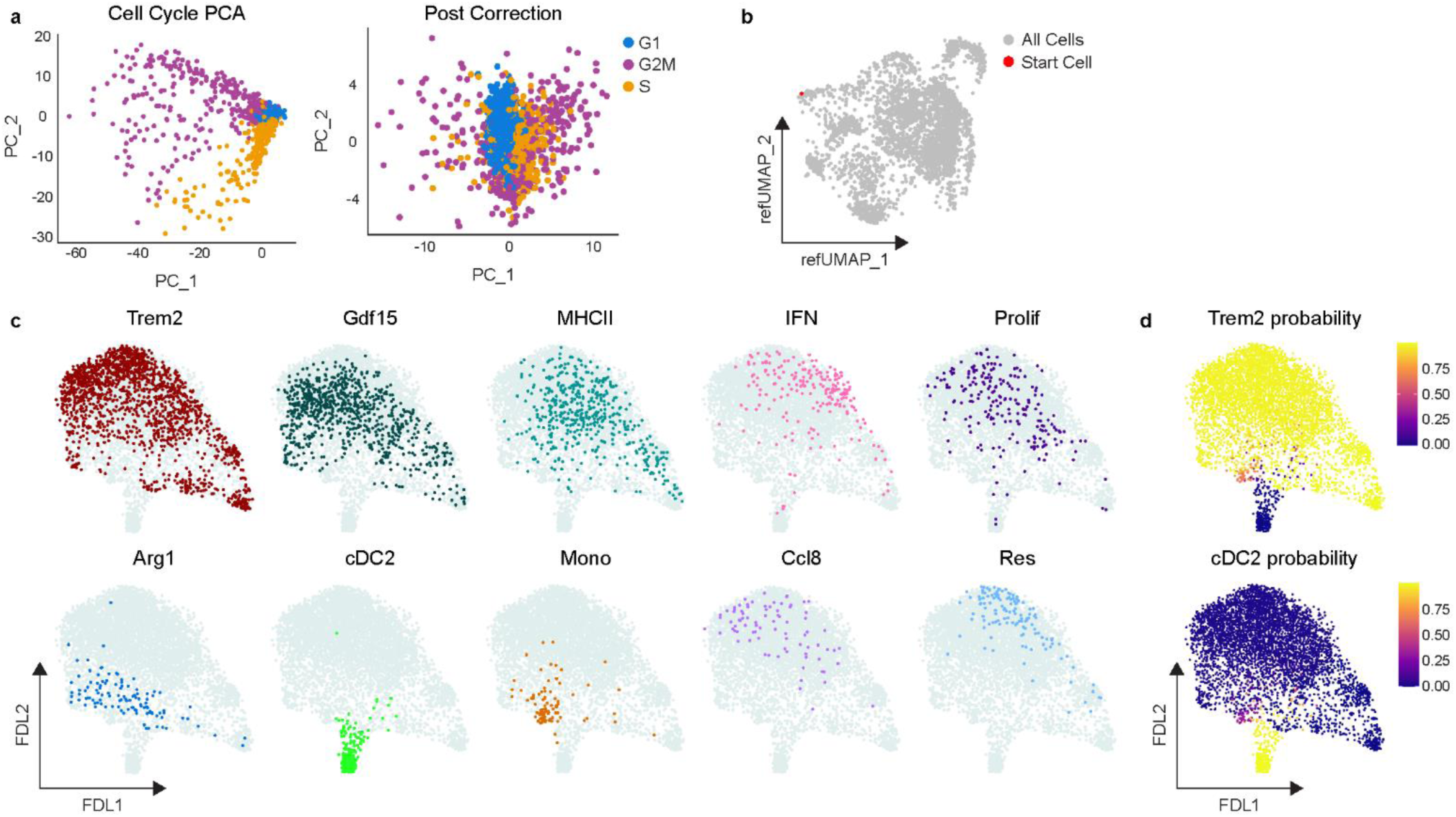
Palantir analysis of Control and macHif1aKO mice 5 days after MI. **a,** PCA plot of cell cycle genes in Control and macHif1aKO scSEQ data 5 days after I/R before (Cell Cycle PCA) and after (Post Correction) regressing out cell cycle scores. **b,** Reference UMAP of Control and macHif1aKO scSEQ data 5 days after I/R highlighting the starting cell used. Starting cell was determined by selecting the Control monocyte with the highest monocyte Z-score (*Plac8*, *Ly6c2*). **c,** Force directed layout (FDL) output of Palantir analysis highlighting each cell subpopulation in Control and macHif1aKO scSEQ data. **d,** Terminal state probability scores from Palantir analysis indicating the likelihood that a cell becomes either Trem2 or cDC2.

**Extended Data Fig. 4.**
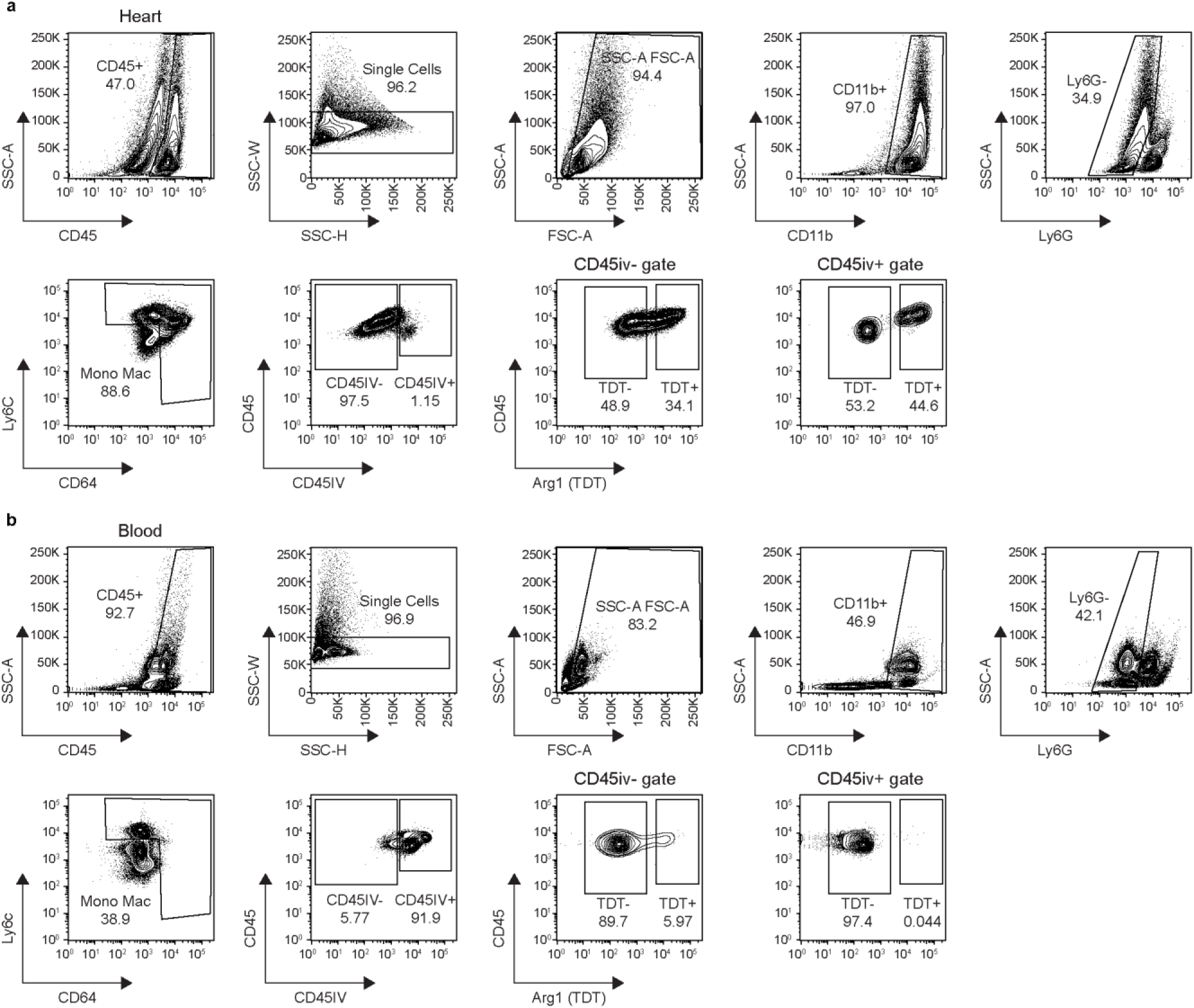
Intravascular staining FACS of *Arg1^tdT-CreERT^*^2^ mice 2 days after MI. **a,** FACS gating strategy for intravascular staining in the heart of *Arg1^tdt-CreERT^*^2^ mice 2 days after I/R. Mice were injected with CD45 antibody 5 minutes prior to tissue collection (CD45IV). Cells were gated for leukocytes (SSC-A vs CD45), single cells (SSC-W vs SSC-H), debris exclusion (SSC-A vs FSC-A), myeloid cells (SSC-A vs CD11b), neutrophil exclusion (SSC-A vs Ly6G) and monocytes/macrophages (Ly6C vs CD64). Intravascular (CD45IV^+^) and extravascular cells (CD45IV^−^) were identified (CD45 vs CD45IV). Arg1^+^ cells were identified via TDT expression in the CD45iv^−^ and CD45iv^+^ gate (CD45 vs Arg1 (TDT)). **b,** Representative FACS plots of the peripheral blood of *Arg1^tdt-CreERT^*^2^ mice 2 days after I/R when performing intravascular staining.

**Extended Data Fig. 5.**
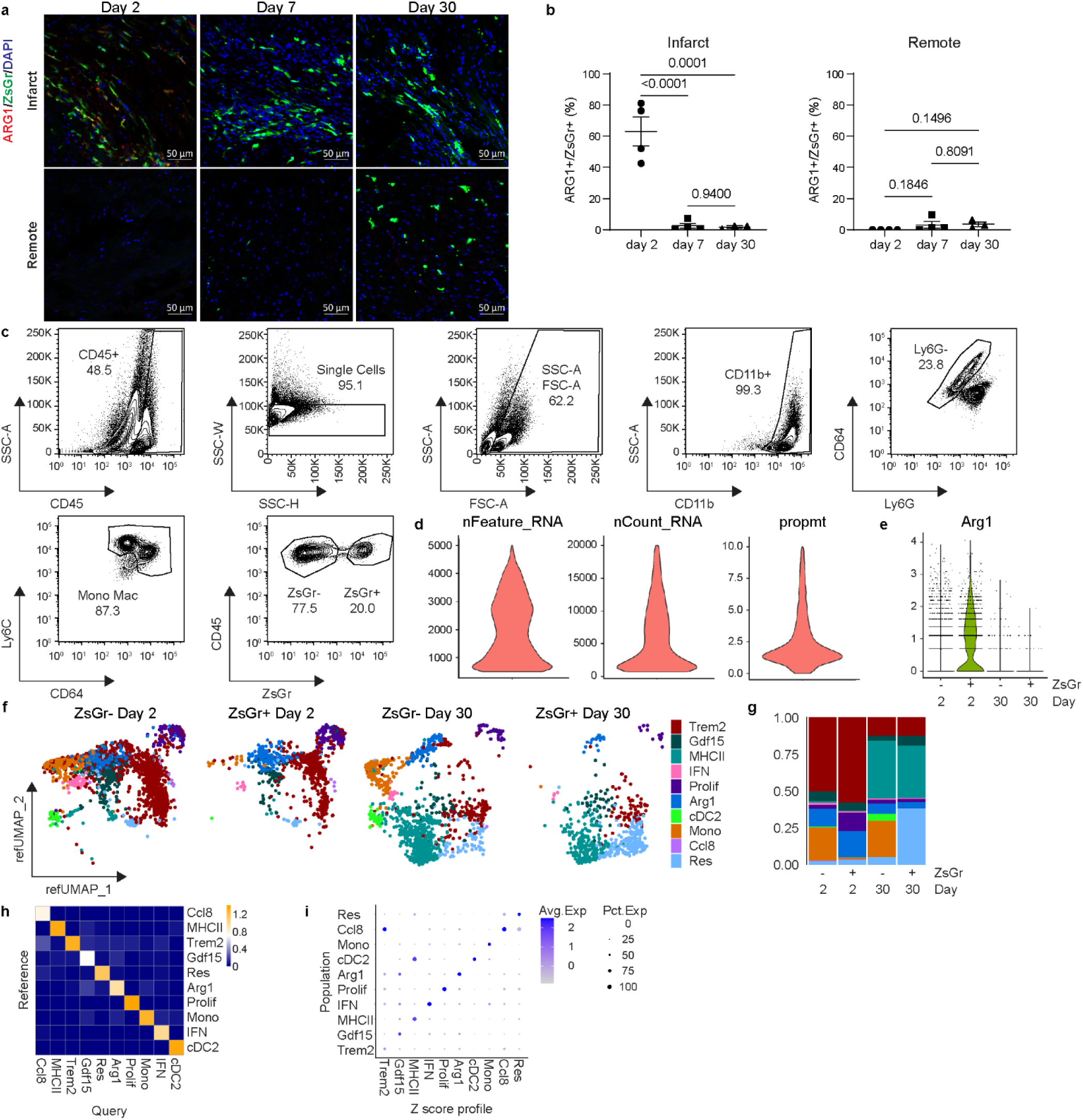
Validation, single cell RNA sequencing, and reference mapping of Arg1ZsGr mice after MI. **a,** Representative 20x confocal images of ARG1 IHC in the infarct and remote zone of Arg1ZsGr mice 2, 7, 30 days after I/R. ARG1 staining is in red, endogenous ZsGr is in green, DAPI is in blue. **b,** Quantification of IHC in a, displayed as he percentage of ZsGr^+^ cells that are ARG1^+^ZsGr^+^. **c,** FACS gating strategy for isolation of ZsGr^−^ and ZsGr^+^ monocytes and macrophages from Arg1ZsGr mice 2 and 30 days after I/R. **d,** Violin plots representing scSEQ data post quality control. Cells with greater than 500 and less than 5000 genes (nFeature_RNA), read counts less than 20,000 (nCount_RNA), and proportion of transcripts mapping to mitochondrial genes less than 10 percent (prompt) were used. **e,** *Arg1* expression in the ZsGr^−^ and ZsGr^+^ libraries at day 2 and 30. **f,** Reference UMAP plots split between ZsGr^−^ and ZsGr^+^ libraries 2 and 30 days after I/R after mapping to reference MI data set. The reference MI scSEQ data set was the same as the one displayed in Extended Data Fig.2. **g,** Stacked bar graph representing the proportion of each cell population in the data split between ZsGr^−^ and ZsGr^+^ cells 2 and 30 days after I/R. **h,** Heat map of mapping scores for each cell population in the Query (Arg1ZsGr) plotted against the Reference MI data set. **i,** Z-score profiles from the reference data set (Extended Data Fig.2f) compared to each subpopulation in the Arg1ZsGr data set represented as a dot-plot. P-values are determined using the ordinary one-way ANOVA for multiple comparisons.

**Extended Data Fig. 6.**
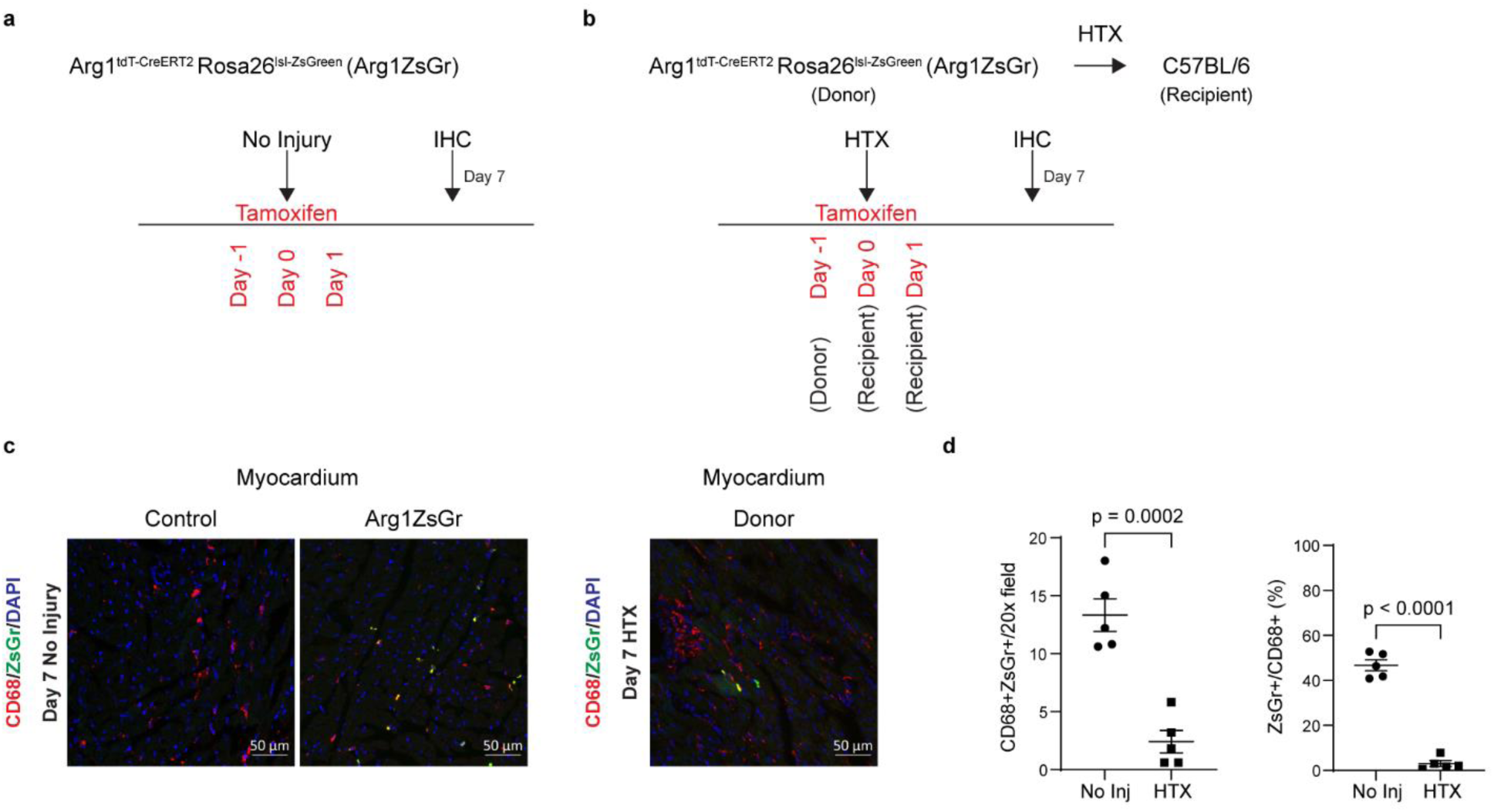
Validation of Arg1ZsGr mice in Injury and Non-Injury conditions. **a,** Schematic of no injury in Arg1ZsGr mice. 60 mg/kg of tamoxifen was injected IP and tissues were collected for IHC Day 7 after not inducing injury. **b,** Schematic of Arg1ZsGr donor heart syngeneic transplant (HTX) into C57BL/6 mice. Arg1ZsGr donor mice were given 60 mg/kg tamoxifen via gavage 1 day prior to transplant. C57BL/6 recipient mice were given 60 mg/kg tamoxifen via gavage 0 and +1 days relative to HTX. Donor hearts were collected 7 days after HTX for IHC. **c,** Representative 20x confocal images of CD68 (red) IHC staining in Control, uninjured Arg1ZsGr, and donor Arg1ZsGr myocardium. Endogenous ZsGr is in green, and DAPI is in blue. **d,** Quantification of IHC in c. Data is displayed as total CD68^+^ZsGr^+^ cells per 20x field, and the percentage of CD68^+^ cells that are CD68^+^ZsGr^+^.

**Extended Data Fig. 7.**
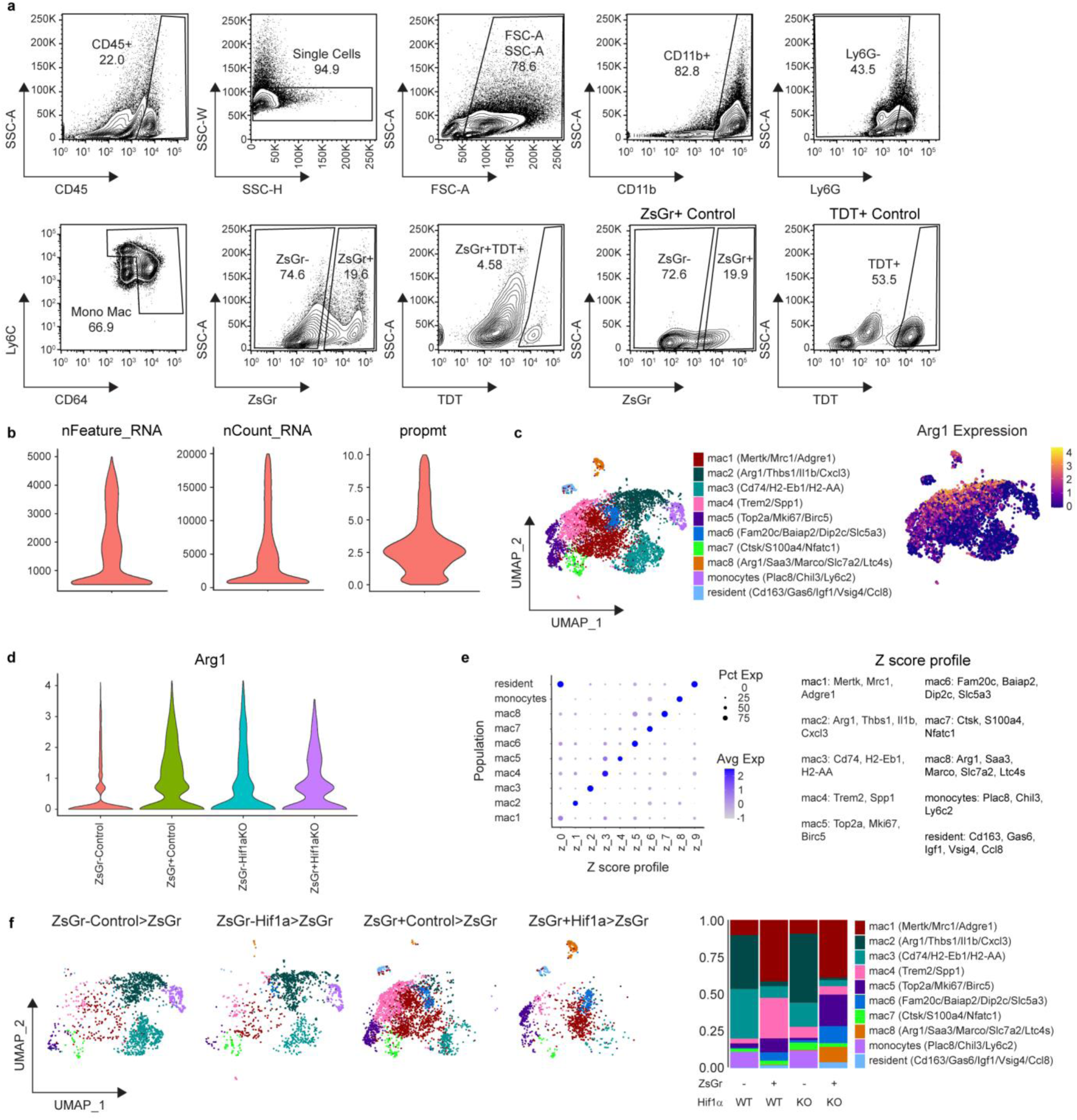
Single cell RNA sequencing of ZsGr- and ZsGr+ cells in Control>ZsGr and Hif1a>ZsGr syngeneic heart transplant mice. **a,** FACS gating strategy for isolation of ZsGr^−^ and ZsGr^+^ monocytes and macrophages from Control>ZsGr and Hif1a>ZsGr mice 5 days after syngeneic heart transplant (HTX) for use in scSEQ. **b,** Violin plots representing scSEQ data post quality control. Cells with greater than 500 and less than 5000 genes (nFeature_RNA), read counts greater than 500 and less than 20,000 (nCount_RNA), and proportion of transcripts mapping to mitochondrial genes less than 10 percent (prompt) were used for downstream analysis. **c,** Annotated UMAP of ZsGr^−^ and ZsGr^+^ cells in Control>ZsGr and Hif1a>Zsgr conditions 5 days after HTX. Arg1 expression plotted on a UMAP projection. **d,** Violin plot of *Arg1* expression split between ZsGr^−^ and ZsGr^+^ cells in Control>ZsGr and Hifa1>ZsGr mice. **e,** Z-score profiles from of each subpopulation in the Control>ZsGr and Hifa1>ZsGr data set represented as a dot-plot. Z-score profile genes for each subpopulation are listed. **f,** Annotated UMAP of scSEQ data 5 days after HTX split between ZsGr^−^ and ZsGr^+^ cells in Control>ZsGr and Hifa1>ZsGr mice, and stacked bar graph representing the proportion of each cell population in the data set split between conditions.

**Extended Data Fig. 8.**
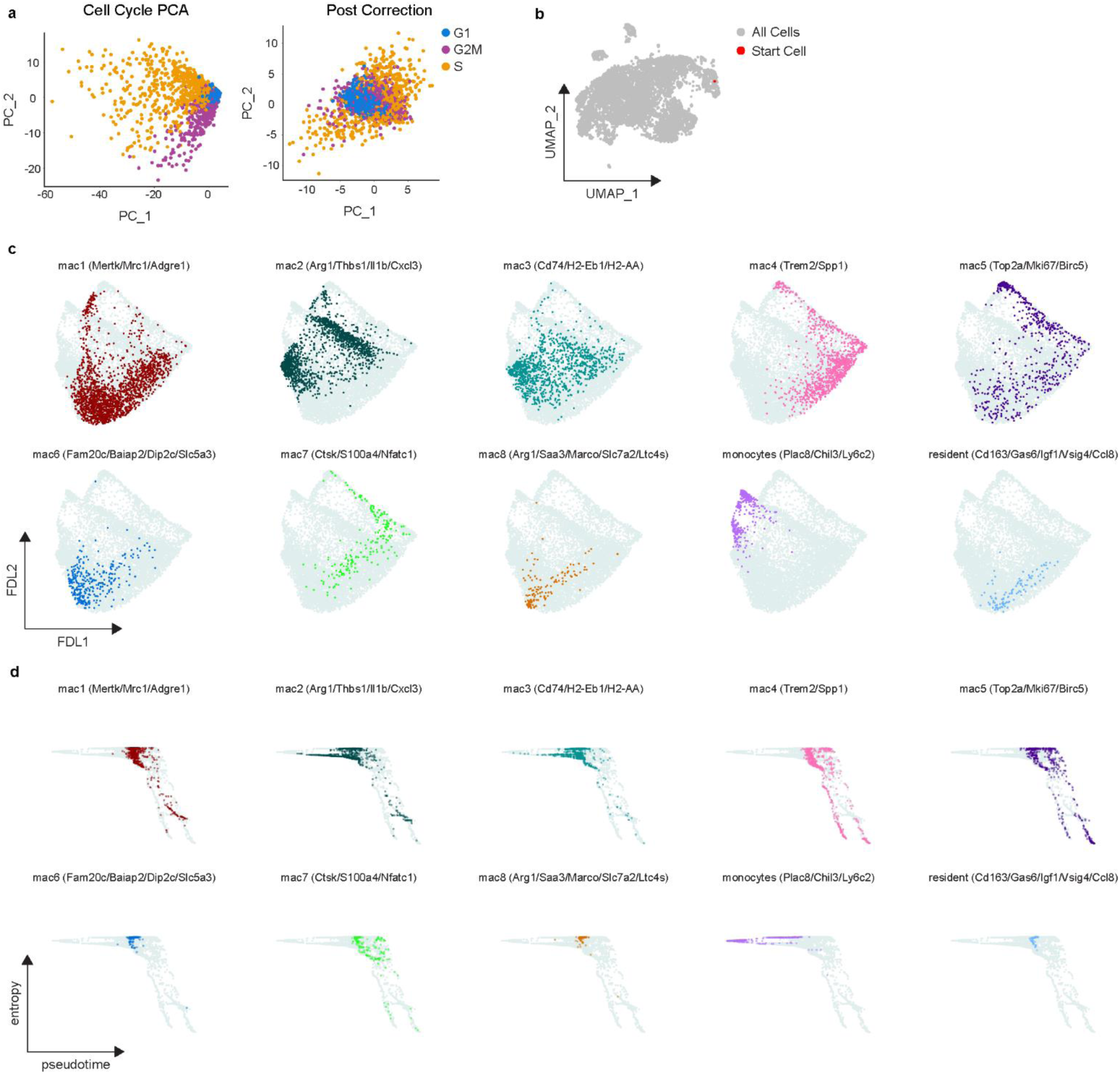
Palantir analysis of ZsGr- and ZsGr+ cells in Control>ZsGr and Hif1a>ZsGr syngeneic heart transplant mice. **a,** PCA plot of cell cycle genes in ZsGr^−^ and ZsGr^+^ cells in Control>ZsGr and Hif1a>ZsGr 5 days after syngeneic heart transplant (HTX) before (Cell Cycle PCA) and after (Post Correction) regressing out cell cycle scores. **b,** UMAP projection of scSEQ data 5 days after HTX highlighting the starting cell used. Starting cell was determined by selecting the monocyte with the highest monocyte Z-score (*Plac8*, *Ly6c2*). **c,** Force directed layout (FDL) output of Palantir analysis highlighting each cell subpopulation in the ZsGr^−^ and ZsGr^+^ cells in Control>ZsGr and Hif1a>ZsGr 5 days after HTX. **d,** Plots of entropy vs pseudotime scores from Palantir analysis for every cell subpopulation in the scSEQ data set.

