## supplementary figures for "Hypoxia Sensing in Resident Cardiac Macrophages Regulates Monocyte Fate Specification following Ischemic Heart Injury"

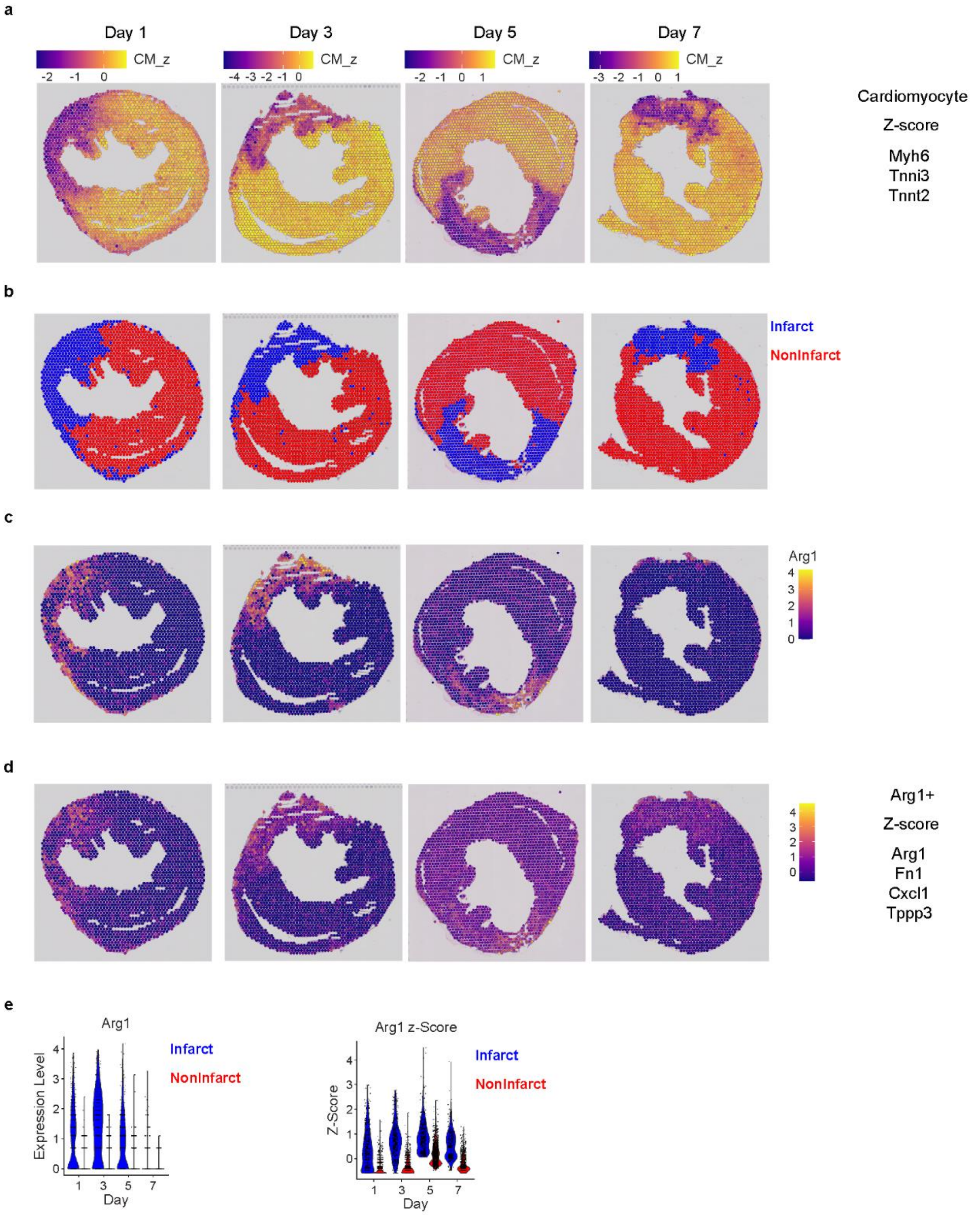


**Supplementary Figure 1. Spatial transcriptomics analysis of *Arg1* in infarcted and non-infarcted regions after MI. a,** Cardiomyocyte gene expression profile (*Myh6*, *Tnni3*, *Tnnt2*) displayed as a Z-score. **b,** Infarct zones were determined as spots that had less than -0.25 cardiomyocyte Z-score. **c,** RNA expression of *Arg1*. **d,** Gene signature of Arg1 macrophages from macHif1aKO data and reference MI dataset (*Arg1*, *Fn1*, *Cxcl1*, *Tppp3*) displayed as a Z-score. **e,** *Arg1* RNA expression and Arg1 macrophage Z-score in infarct (blue) and non-infarct (red) regions displayed as a violin plot.

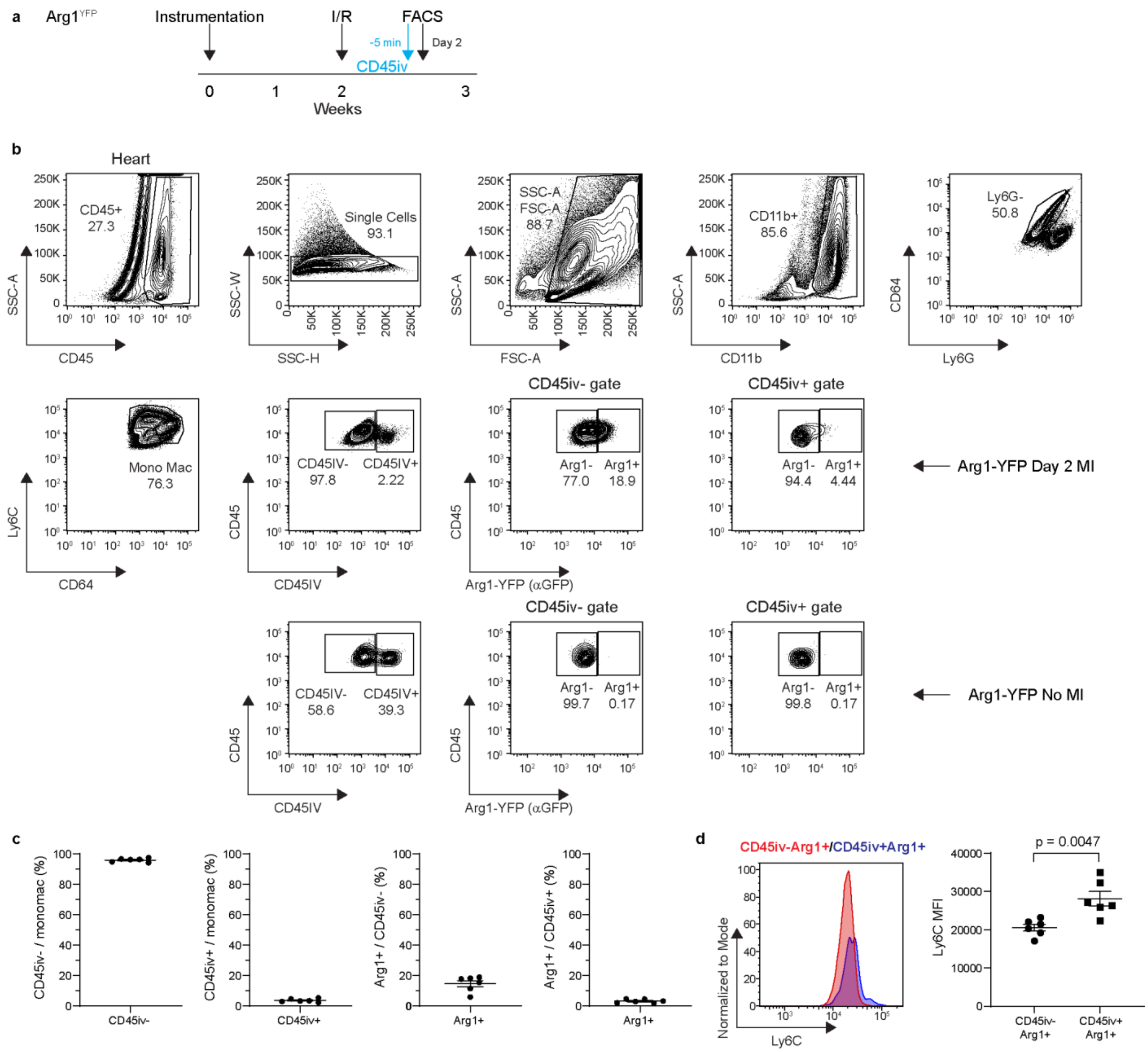


**Supplementary Figure 2. Intravascular staining FACS of *Arg1*$^{YFP}$ mice 2 days after MI a,** Schematic of intravascular FACS in *Arg1*$^{YFP}$ mice 2 days after I/R. Mice were injected with a CD45 antibody 5 minutes prior to tissue collection. **b,** FACS gating strategy for intravascular staining in the heart of *Arg*$^{YFP}$ mice 2 days after I/R or no injury. Cells were gated for leukocytes (SSC-A vs CD45), single cells (SSC-W vs SSC-H), debris exclusion (SSC-A vs FSC-A), myeloid cells (SSC-A vs CD11b), neutrophil exclusion (CD64 vs Ly6G) and monocytes/macrophages (Ly6C vs CD64). Intravascular (CD45IV+) and extravascular cells (CD45IV-) were identified (CD45 vs CD45IV). Arg1$^+$ cells were identified via intracellular anti-GFP staining in the CD45iv- and CD45iv+ gate (CD45 vs Arg1-YFP (αGFP)). **c,** Quantification of populations of interest from Day 2 MI mice in b. **d,** Representative histogram of Ly6C staining between CD45iv$^-$Arg1$^+$ cells and CD45iv$^+$Arg1$^+$ cells. Quantification of Ly6C mean fluorescence intensity (MFI) between CD45iv$^-$Arg1$^+$ cells and CD45iv$^+$Arg1$^+$ cells. P-values are determined using a two tailed t-test assuming equal variance.

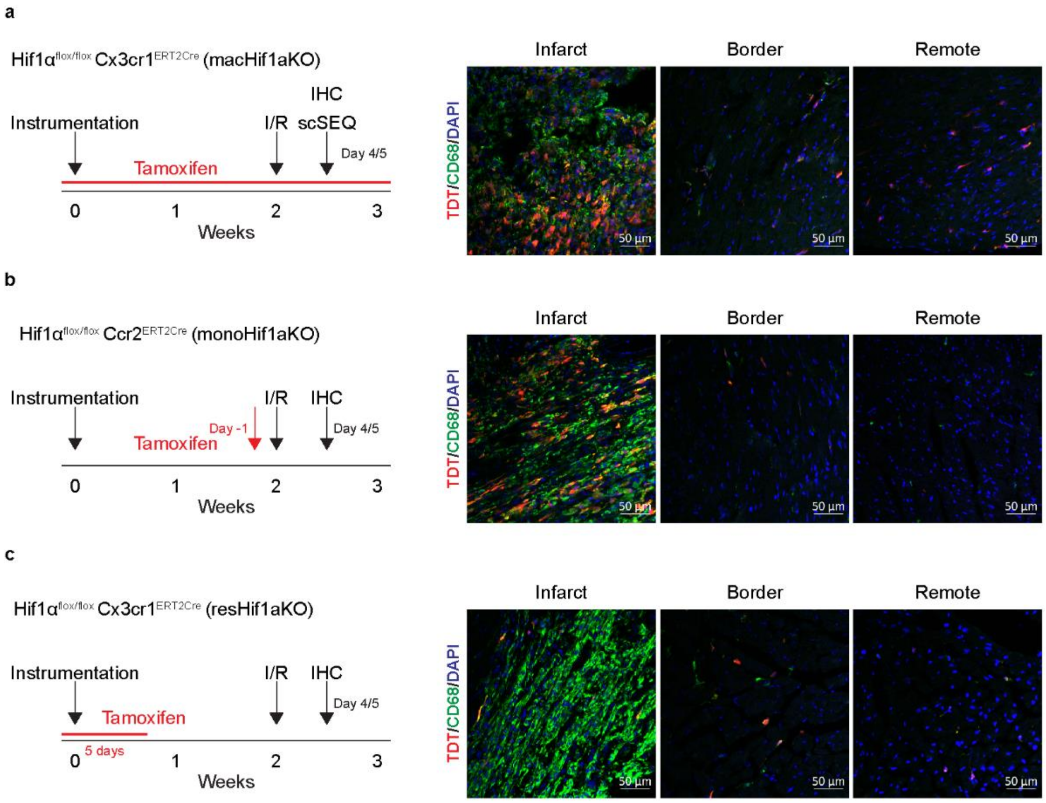


**Supplementary Figure 3. Tamoxifen strategies to induce Cre recombination in all monocytes and macrophages, recruited monocytes and macrophages, or cardiac resident macrophages. a,** Schematic of experimental design and representative 20x confocal images in the infarct, border, and remote zone when *Hif1α* is knocked out in all monocytes and macrophages during closed chest I/R (macHif1aKO). **b,** Schematic of experimental design and representative 20x confocal images in the infarct, border, and remote zone when *Hif1α* is knocked out in monocytes and monocyte derived macrophages during closed chest I/R (monoHif1aKO). **c,** Schematic of experimental design and representative 20x confocal images in the infarct, border, and remote zone when *Hif1α* is knocked out in resident macrophages during closed chest I/R (resHif1aKO). Endogenous TDT signal is in red, CD68 IHC staining is in green, and DAPI is in blue.

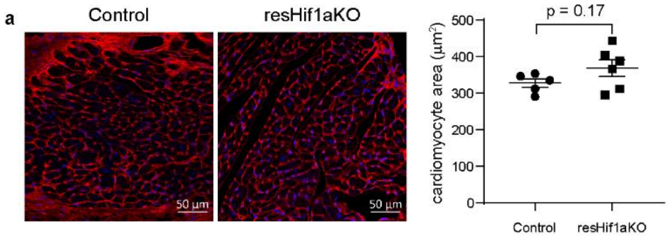


**Supplementary Figure 4. WGA staining in resHif1aKO mice a,** Representative WGA staining of the border zone of the infarct of Control and resHif1aKO hearts 4 weeks after I/R. Quantification of the average cardiomyocyte area was determined from WGA staining. P-values are determined using a two tailed t-test assuming equal variance.

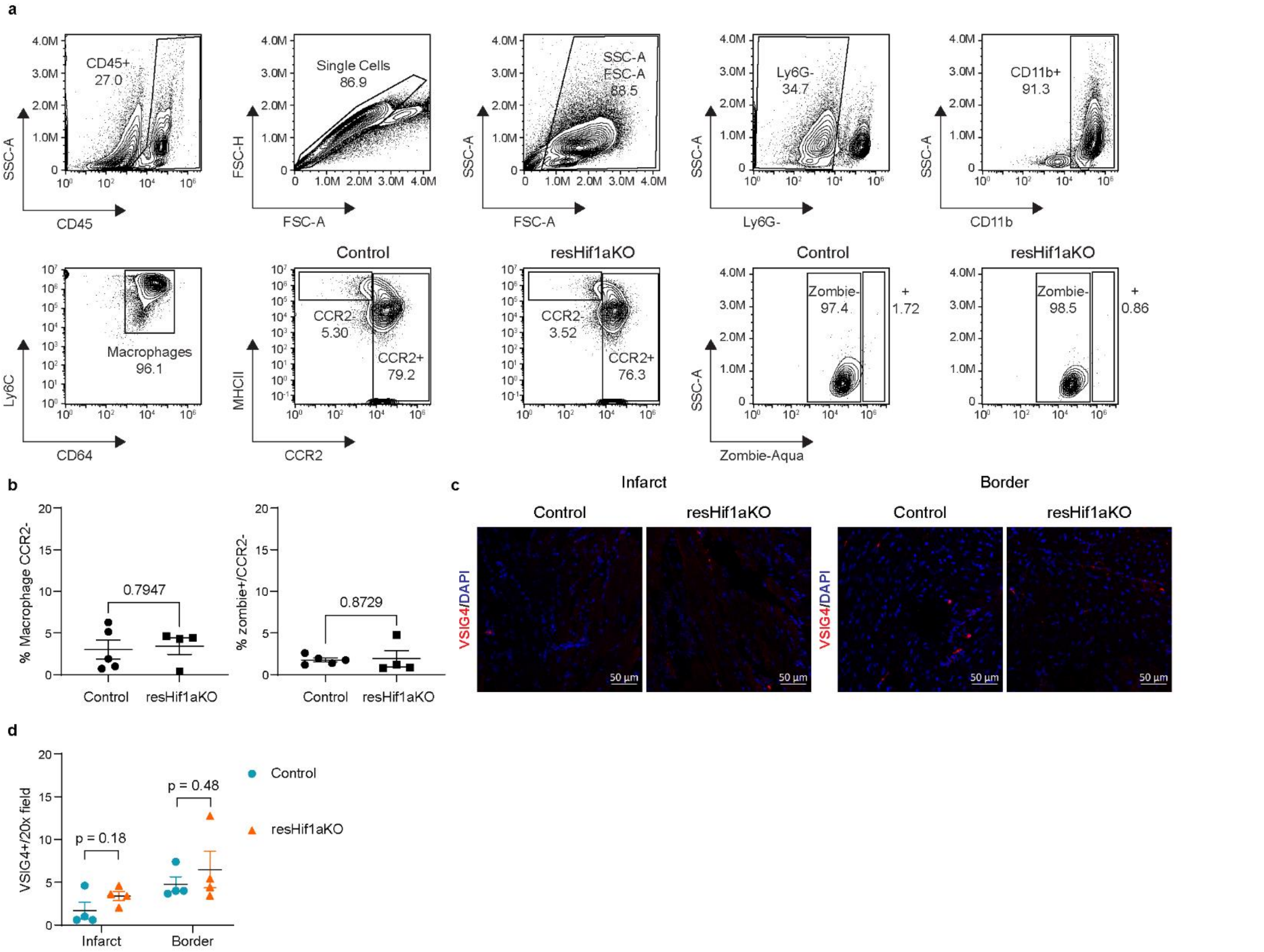


**Supplementary Figure 5. *Hif1α* deletion in resHif1aKO mice does not affect resident cardiac macrophage cell death after MI. a,** FACS gating strategy for resHif1aKO mice 1 day after I/R. Cells were gated for leukocytes (SSC-A vs CD45), single cells (FSC-H vs FSC-A), debris exclusion (SSC-A vs FSC-A), neutrophil exclusion (SSC-A vs Ly6G), myeloid cells (SSC-A vs CD11b), macrophages (Ly6C vs CD64), and resident ($CCR2^-$) vs recruited ($CCR2^+$) macrophages (MHCII vs CCR2). Cell death rate in $CCR2^-$ macrophages was determined by Zombie staining (SSC-A vs Zombie-Aqua). **b,** Quantification of cell types of interests in a. The percentage of macrophages that are $CCR2^-$ is displayed, followed by the percentage of $CCR2^-$ macrophages that are $Zombie^+$. **c,** Representative 20x confocal images of resident macrophage marker VSIG4 (red) and DAPI (blue) IHC in the infarct and border zones of Control and resHif1aKO mice 1 day after I/R. **d,** Quantification of IHC in c, displayed as the total number of VSIG4+ cells per 20x field.
